# Standardizing and applying a mating-based whole-genome simulation approach reveals caution in using chromosome-level PCA and kinship estimates

**DOI:** 10.1101/2023.03.16.532885

**Authors:** Zuxi Cui, Fredrick R. Schumacher

**Author notes:** Corresponding Author: Fredrick R. Schumacher, PhD, MPH, Department of Population and Quantitative Health Sciences School of Medicine, Case Western Reserve University 10900 Euclid Avenue, Cleveland, OH 44106-4945.

## Abstract

This paper presents a new and efficient method for simulating pseudo-genotype data using the standardized protocol of SLiM, which offers a flexible alternative to traditional methods that rely on large genetic datasets. These datasets can be time-consuming to obtain, especially when institutional review board (IRB) review is involved, making simulation an attractive alternative. While HapGen v2 is the most popular genotype simulator, we found that SLiM has the potential for more customizable simulation to meet multiple needs.

To validate our new method, we compared its performance among parallel simulations varying multiple parameters. Our results showed that SLiM is capable of simulating samples up to 333 times the input size, with a low rate of simulated samples that are 2nd or closer relatives (REV), making it a promising alternative to HapGen. We also applied our whole-genome simulation approach to sensitivity analyses of chromosome-level principal component analysis (PCA) and kinship estimation. Our findings revealed important insights into the sensitivity of PCA and kinship estimation, highlighting the unequal distribution of population structure across chromosomes and ancestries. Furthermore, our study provides experimental support for avoiding chromosome-level quality control statistics.

Overall, our standardized protocol of SLiM offers a flexible new way to produce pseudo-genotype data, and our findings provide valuable insights that can advance research in the field. By demonstrating the potential of SLiM for more customizable simulations and highlighting the importance of considering the distribution of population structure across chromosomes and ancestries, our research has significant implications for the study of genetics and genomics.

**Author Summary:** In this publication, we introduce a novel approach to genotype simulation using a mating-based strategy in SLiM. Our approach mimics mitosis computationally and stands out as the only one available as of December 2022 that can maintain cross-chromosome associations during whole-genome level simulation, with no competitors in sight. Additionally, our approach is applicable to regional or chromosomal genotype simulation. When compared to the current gold-standard chromosome-level simulator, HapGen, our approach exhibits superior performance when generating large sample sizes (>13,000). We provide an application example that uses whole-genome simulation to underscore the importance of whole-genome quality control (QC) statistics, such as principal component analysis (PCA) and kinship estimates, compared to the chromosome-level ones. Results of the application indicate instability and bias in the chromosome-level QC statistics. Overall, our approach represents a valuable tool for genetics research that can assist in the evaluation and validation of genetic analyses, and people should avoid chromosome-level QC statistics.

## Introduction

The recent influx of whole-genomic data has fostered the development of novel statistical methods for the analysis and discovery of gene-phenotype associations. However, access to a comprehensive whole-genome reference dataset used for better understanding the parameters of these novel statistical methods is crucial. While the cost of sequencing continues to decrease, access to real data is limited, as most whole-genome sequencing data is either private or controlled-accessed. Furthermore, publicly available genotype datasets consist mainly of individuals of European continental ancestry, such as the Haplotype Reference Consortium (HRC) (1). The majority of HRC’s haplotype reference panel is not publicly available, and over 95% of the samples are of European ancestry. In addition, data use agreements and institutional review board approval further restrict access to this data. One alternative to address this issue is to use genotype simulation tools to create pseudo-genotypes for validating statistical methods.

One widely used genotype simulator is HapGen v2, which has been cited over 300 times as of June 2022 (2). HapGen employs the “product of approximate conditionals” (PAC) model to relate linkage disequilibrium (LD) patterns to the underlying recombination rate (3). This model allows the preservation of allele frequencies and LD structure from the reference panel. However, the resampling method used by HapGen to generate diversity can produce an overrepresentation of increased relatedness in situations requiring a high simulation ratio between output and input sample sizes. Furthermore, HapGen is limited to chromosome-level simulations only.

In contrast, SLiM v3.3, released by the Messer’s lab in late 2019, is the first whole-genome simulator that uses a forward-time Non-Wright-Fisher nucleotide-based model to create offspring of a reference panel in the popular variants calling format (VCF) (4,5). This model allows individuals to undergo a dynamic process with new offspring add-ins from random bi-parental mating and age-sensitive kick-outs. Mating and death patterns apply to all chromosomes, providing a solution to whole-genome data simulation maintaining cross-chromosome associations. SLiM also allows the setting of variable recombination sites within each simulation. However, there is a lack of information on best practices for using SLiM to create genome-wide pseudo data. Thus, we conducted this study to compare SLiM with HapGen regarding relativeness and ancestral accuracy, and to provide guidelines for preferred scenarios.

We ran parallel simulations varying several parameters to identify best practices for using SLiM. Because HapGen does not support whole-genome simulation, all QC statistics were derived at the chromosome level and inter-software comparisons were by chromosomes. However, recent evidence showed a potential that the effect of population structure differed along the genome, so, as an application, a whole-genome simulation was performed to explore the effect modification of population structure by chromosomes (6). In the application, we tested whether the sensitivities of chromosome-level QC statistics, such as principal component analysis and kinship estimation, vary by chromosomes.

## Objectives

The main objective of our study was to provide a standardized solution for creating high-quality whole-genome-level pseudo genotype data to increase the sample size of open-access data. We conducted sensitivity analyses of controllable parameters to determine their impact on quality scenarios and optimized sets. As a secondary objective, we evaluated the sensitivity of chromosome-level quality control (QC) statistics using our whole-genome simulation approach.

## Results

### Reference QC

#### Reference for simulation

The exclusion of variants during each QC step varied across the different subpopulations (Figure 3). The 1000 Genomes Project dataset comprised 2,504 samples from diverse populations and included over 81.3 million variants. Upon ancestry stratification, we obtained 13.0 million variants in 103 CHB samples, 12.9 million variants in 91 GBR samples, and 22.4 million variants in 108 YRI samples. Following the exclusion of INDELs, approximately 10% of variants were lost, resulting in a final count of 11.8 million, 11.7 million, and 20.5 million variants for CHB, GBR, and YRI, respectively. We identified 38 tri-allelic loci in chromosomes 3 and 8, which were subsequently excluded. Additionally, we removed one sample (NA19240) from family Y117 as she was the offspring of samples NA19238 and NA19239. To cover different numbers of variants (ranging from 8.5 million to 19.6 million) across different ancestries and sample sizes (30, 60, and 90), we generated nine subsets using random sampling.

**Figure 1:**
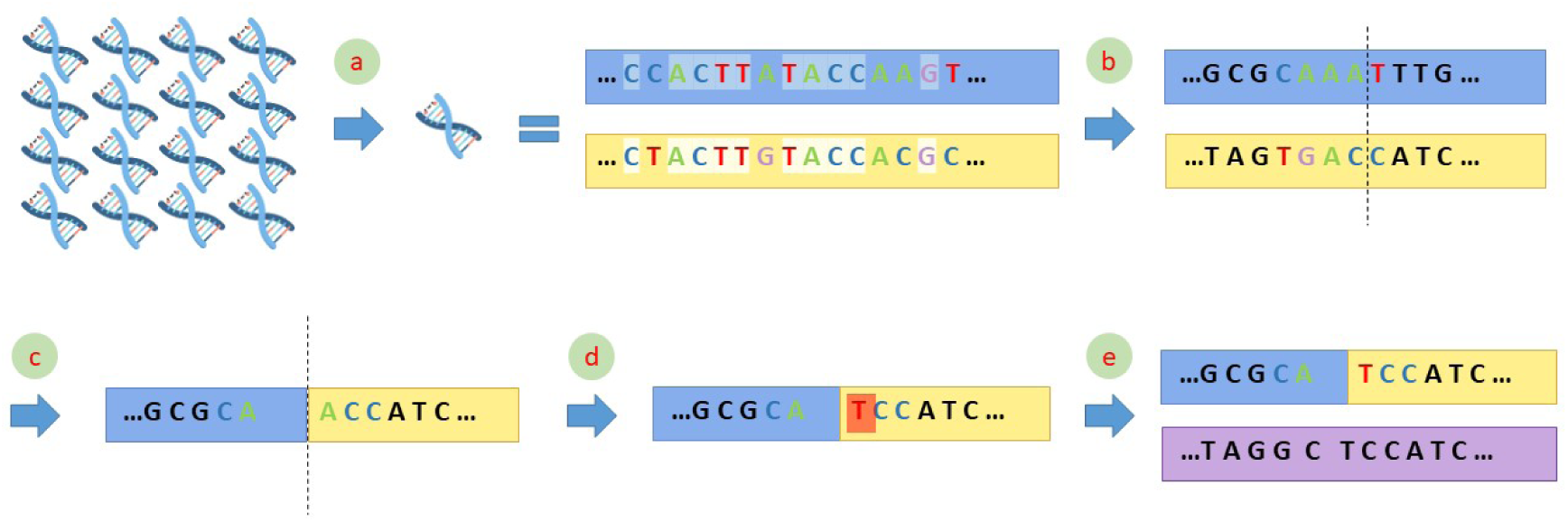
Simulation process of one generation pass. One generation pass includes five steps. **a**. Chromosome-level random sampling. Each symbol represents a sequence of the same chromosome from different people. Random sampling applies. **b**. Non-variant exclusion. Shadow marks non-variant sites to be excluded in the QC. **c**. Recombination. Dash line marks where recombination occurs. **d**. Mutation. Orange shadow marks mutation site. Both recombination and mutation sites can be customized via modeling. Uniform distribution applied in our study. **e**. Pairing. Purple chromosome is from another sample after the same process of 1-4. Combination of two chromosomes makes a new offspring.

**Figure 2:**
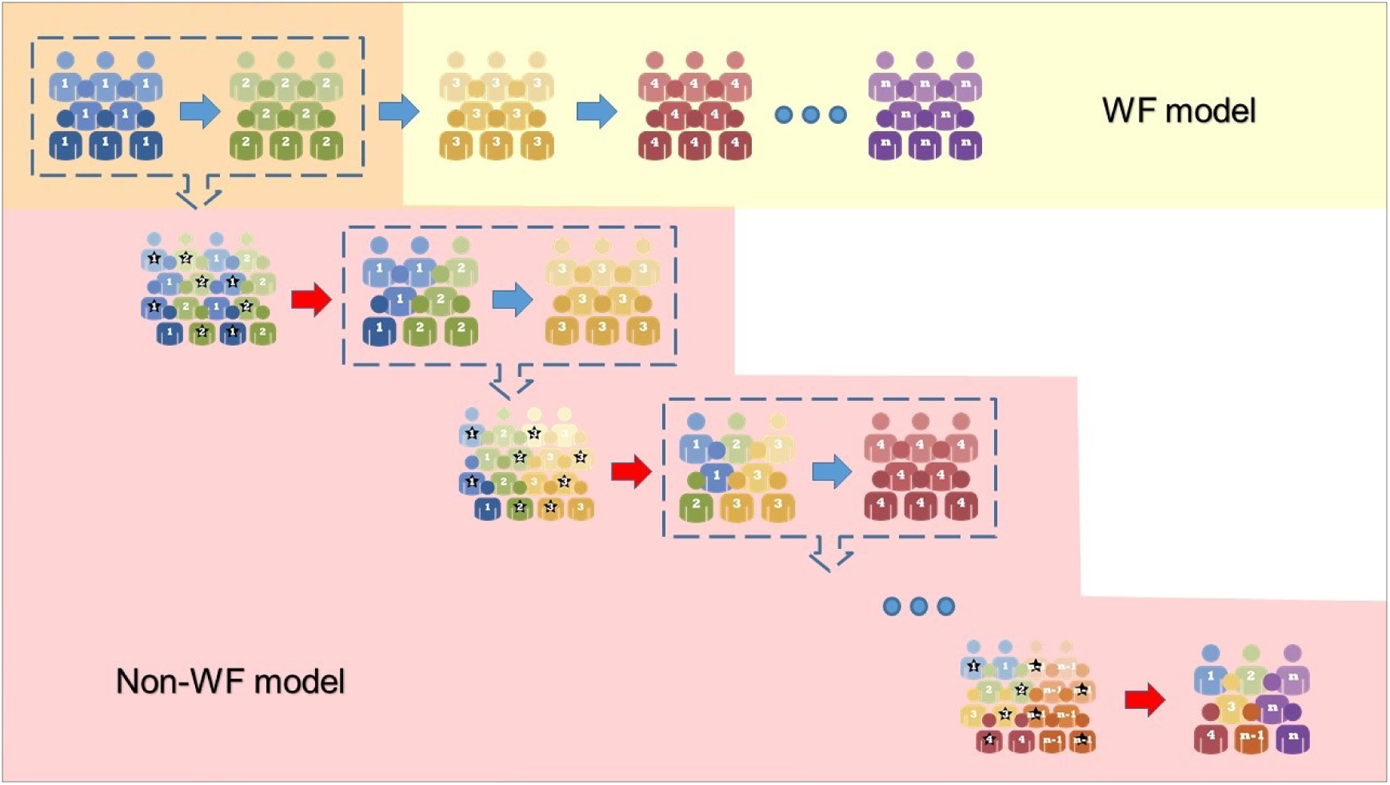
Workflow of the third stage in WF and Non-WF model. Difference between third stage in WF and Non-WF model. Corn silk marks the third stage in the WF model and dusty rose marks that in the Non-WF model. Founders are dropped after mating at each generation pass in the WF model but pooled with offspring for a sampling process in the Non-WF model. Blue arrow represents creating a new generation by mating. Red arrow represents a sampling procedure keeping random selected samples with a star mark. Dash rectangle represents a pooling process under a fitness model.

**Figure 3:**
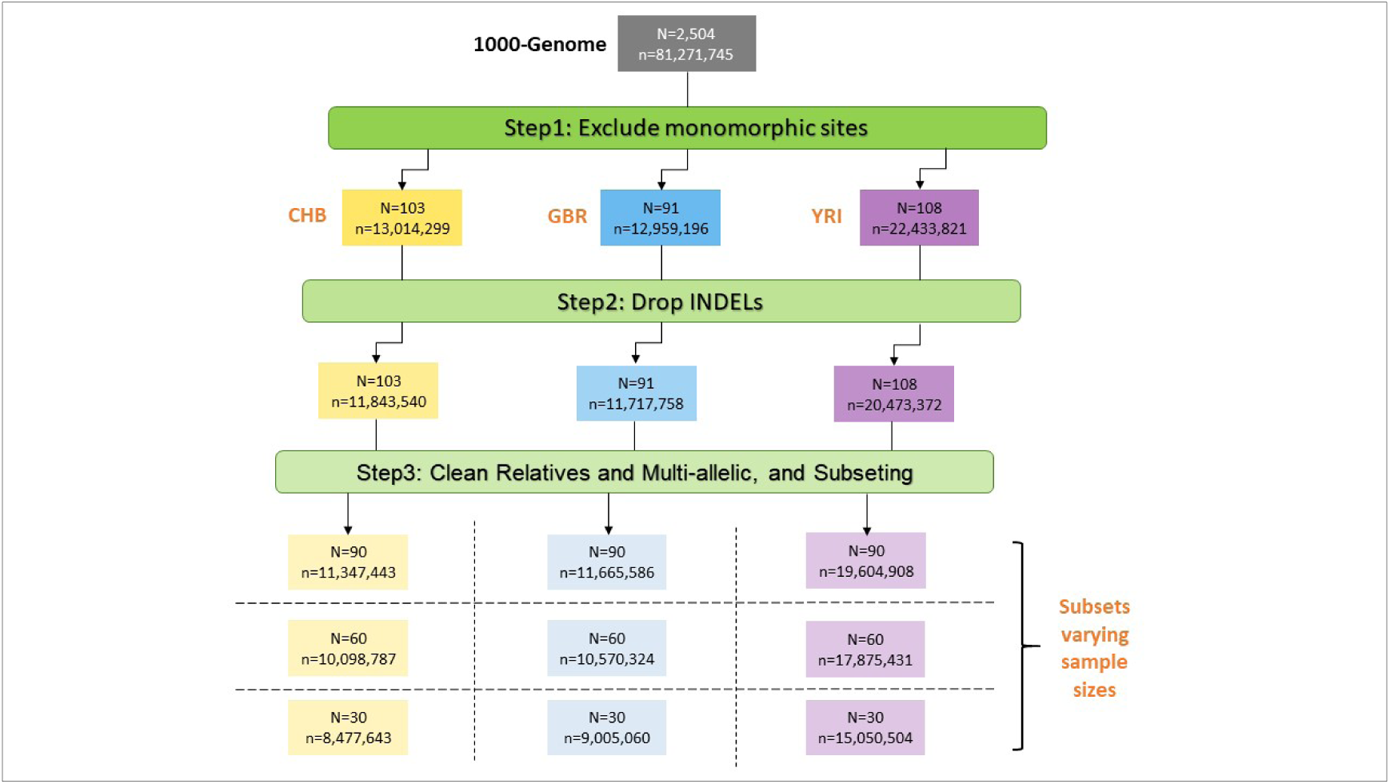
Summary of sample sizes and number of variants after each step of Pre-simulation QC. N: the sample size; n: the total number of variants.

The post-QC reference datasets showed that the distribution of MAFs varied inversely with the number of variants in each MAF chunk, as depicted in Figure S5. The distribution patterns were comparable across the three subpopulations. The distributions for CHB and GBR were similar, with a 30-sample increase resulting in approximately 1.5M more rare variants (MAF < 0.01). In contrast, the YRI reference showed twice the difference in the number of rare variants (~3M) when the sample size was increased by 30.

#### Reference for PCA and kinship analysis

After the pruning and exclusion of uncommon variants, the QC reference array was reduced from 81 million to 242 thousand variants. The number of variants that were retained varied depending on the chromosome, with smaller chromosomes having fewer variants (Figure S6). Specifically, chromosome one had the largest number of variants retained, with 19,573 variants, while chromosome 22 had the smallest number, with only 4,243 variants.

### Simulated variants

According to Table 1, mean number of simulated variants across ten parallel simulations indicated a decline due to generations of mating. The loss of variants ranged from 9.1% to 75.1% of those in the reference panel, upon varying parameters. CHB with 30 input and 30 generation passes from Non-WF model experienced a minimal loss of 77K (9.1%) variants, while YRI with 90 input and 300 generation passes from Non-WF model lost the maximum of 14.7M (75.1%) variants. Interestingly, while YRI references covered most variants, they experienced the most significant loss of variants both identical and percentage-wise during simulations, followed by CHB and GBR. The loss became more pronounced with an increase in the number of generation passes. Cumulatively, the reference data, which included 90 YRI and 19.6M variants, lost 6.7M (34.4%), 10.7M (54.7%), and 14.3M (72.8%) after 30, 100, and 300 generations, respectively, under the WF model. Notably, the rate of variant loss slowed as the number of generations increased. The number of variants lost from 0-30 generations was roughly twice the number lost from 30-100 or 100-300 generations (6.7M vs. 4M/3.6M). Similar trends in variant loss were observed in simulations under the Non-WF model.

**Table 1:** Number of Post-simulation variants averaged by ten parallel simulations

|  | N input | 30 |  |  | 60 |  |  | 90 |  |  |
| --- | --- | --- | --- | --- | --- | --- | --- | --- | --- | --- |
|  | n gen runs | 30 | 100 | 300 | 30 | 100 | 300 | 30 | 100 | 300 |
| WF model | Subpopulation=YRI |  |  |  |  |  |  |  |  |  |
|  | Input variants | 15,050,504 |  |  | 17,875,431 |  |  | 19,604,908 |  |  |
|  | Total variants output | 12,190,856 | 8,638,932 | 5,266,252 | 12,681,907 | 8,829,647 | 5,283,793 | 12,856,114 | 8,885,735 | 5,339,266 |
|  | Variant loss | 2,859,648 | 6,411,572 | 9,784,252 | 5,193,524 | 9,045,784 | 12,591,638 | 6,748,794 | 10,719,173 | 14,265,642 |
|  | %loss | **19.0%** | **42.6%** | **65.0%** | **29.1%** | **50.6%** | **70.4%** | **34.4%** | **54.7%** | **72.8%** |
|  | Subpopulation=CHB |  |  |  |  |  |  |  |  |  |
|  | Input variants | 8,477,643 |  |  | 10,098,787 |  |  | 11,347,443 |  |  |
|  | Total variants output | 7,109,817 | 5,513,093 | 3,684,432 | 7,333,632 | 5,598,556 | 3,693,818 | 7,428,685 | 5,648,134 | 3,687,949 |
|  | Variant loss | 1,367,826 | 2,964,550 | 4,793,211 | 2,765,155 | 4,500,231 | 6,404,969 | 3,918,758 | 5,699,309 | 7,659,494 |
|  | %loss | **16.1%** | **35.0%** | **56.5%** | **27.4%** | **44.6%** | **63.4%** | **34.5%** | **50.2%** | **67.5%** |
|  | Subpopulation=GBR |  |  |  |  |  |  |  |  |  |
|  | Input variants | 9,005,060 |  |  | 10,570,324 |  |  | 11,665,586 |  |  |
|  | Total variants output | 7,619,658 | 5,944,344 | 3,879,731 | 7,914,601 | 6,008,754 | 3,931,173 | 8,077,362 | 6,064,149 | 3,932,165 |
|  | Variant loss | 1,385,402 | 3,060,716 | 5,125,329 | 2,655,723 | 4,561,570 | 6,639,151 | 3,588,224 | 5,601,437 | 7,733,421 |
|  | %loss | **15.4%** | **34.0%** | **56.9%** | **25.1%** | **43.2%** | **62.8%** | **30.8%** | **48.0%** | **66.3%** |
| Non-WF model | Subpopulation=YRI |  |  |  |  |  |  |  |  |  |
|  | Input variants | 15,050,504 |  |  | 17,875,431 |  |  | 19,604,908 |  |  |
|  | Total variants output | 13,328,624 | 9,502,403 | 5,263,189 | 13,786,447 | 9,207,230 | 5,358,977 | 13,844,243 | 9,662,147 | 4,878,518 |
|  | Variant loss | 1,721,880 | 5,548,101 | 9,787,315 | 4,088,984 | 8,668,201 | 12,516,454 | 5,760,665 | 9,942,761 | 14,726,390 |
|  | %loss | **11.4%** | **36.9%** | **65.0%** | **22.9%** | **48.5%** | **70.0%** | **29.4%** | **50.7%** | **75.1%** |
|  | Subpopulation=CHB |  |  |  |  |  |  |  |  |  |
|  | Input variants | 8,477,643 |  |  | 10,098,787 |  |  | 11,347,443 |  |  |
|  | Total variants output | 7,708,929 | 5,341,134 | 3,831,047 | 7,224,268 | 5,949,535 | 3,721,613 | 7,950,507 | 5,742,014 | 3,338,627 |
|  | Variant loss | 768,714 | 3,136,509 | 4,646,596 | 2,874,519 | 4,149,252 | 6,377,174 | 3,396,936 | 5,605,429 | 8,008,816 |
|  | %loss | **9.1%** | **37.0%** | **54.8%** | **28.5%** | **41.1%** | **63.1%** | **29.9%** | **49.4%** | **70.6%** |
|  | Subpopulation=GBR |  |  |  |  |  |  |  |  |  |
|  | Input variants | 9,005,060 |  |  | 10,570,324 |  |  | 11,665,586 |  |  |
|  | Total variants output | 7,129,196 | 6,129,693 | 3,783,113 | 7,815,600 | 6,522,978 | 4,146,136 | 7,322,827 | 6,054,631 | 4,100,348 |
|  | Variant loss | 1,875,864 | 2,875,367 | 5,221,947 | 2,754,724 | 4,047,346 | 6,424,188 | 4,342,759 | 5,610,955 | 7,565,238 |
|  | %loss | 20.8% | 31.9% | 58.0% | 26.1% | 38.3% | 60.8% | 37.2% | 48.1% | 64.9% |

Figure 4 displays histograms of MAFs in simulated data under the WF model, where the input size was 90. The histograms revealed that the number of lost variants was inversely related to their MAF. The majority of lost variants were less common (MAF < 0.5) or rare (MAF < 0.1).

**Figure 4:**
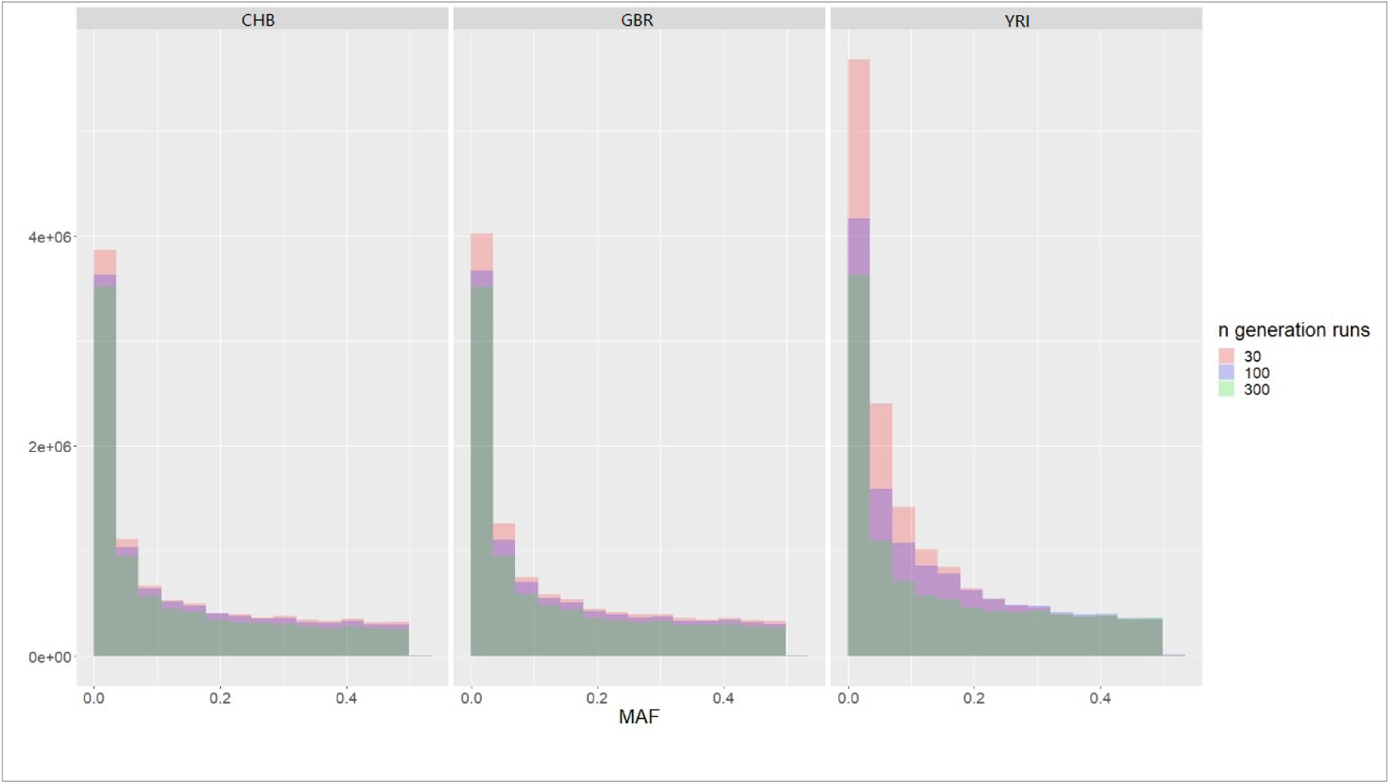
Distribution of MAF of data from simulations varying number of generation passes. Histograms present counts of MAF of data simulated based on an input sample size of 90 under WF model. Light green shadow shows simulations with 300 generation passes. Blue shows simulations with 100 generation passes. Dusty rose shows simulations after 30 generation passes. Dark green shows overlap of three histograms. Purple shows overlap between simulations after 100 and 300 passes.

Although YRI had a significantly larger number of variants, especially rare ones, during the initial mating stages than the other two subpopulations, the gap gradually reduced to almost zero after 300 generation passes.

### Intro-software practice

#### Robustness - Frequencies of nucleotides

Across ten simulations, mean frequencies for adenine (A), cytosine (C), guanine (G), and thymine (T) varied by ancestries and chromosomes, with the maximum standard deviations being 7.90×10-6, 8.22×10-6, 8.07×10-6, and 8.23×10-6 for the four nucleotides. Figure S7 displays boxplots illustrating the largest potential for variation under the assumptions described in the methods. Based on the boxplots, five medians overlapped with the lower quartile, four overlapped with the upper quartile, and three overlapped with both. Over 70% of frequencies were within the boxes, and only one or no frequency in each series fell outside of the 95% confidence interval. The same analyses were conducted on simulations with different parameter settings, revealing smaller variations in nucleotide frequencies within larger chromosomes, with fewer generation passes and larger input sample sizes, and in the Non-WF model.

#### Robustness - PC-dispersion

The PCA plots revealed that the standard deviation of PCs increased with each generation pass, resulting in a wider dispersion of simulated data points (as seen in Figures S8 A-F & S9 A-F). Interestingly, ten parallel simulations with either 30 or 100 generation passes showed little to midcore dispersion and were clustered in the same region of the referenced samples. In contrast, simulations with 300 generation passes displayed a substantial degree of dispersion, indicating greater genetic divergence. We also observed that the degree of spread in PC plots was similar for the three subpopulations (Figures S8&S9: AB vs. CD vs. EF), implying that they had undergone similar levels of genetic drift. Additionally, when other parameters were held constant, there was no significant difference in the robustness of the degree of dispersion between the WF and Non-WF models (Figures S8&S9: A vs. B, C vs. D, E vs. F). Furthermore, we found that the dispersion of PC plots was more pronounced in chromosome 22 simulations compared to chromosome one simulations. This suggests that smaller chromosomes may be more prone to genetic drift than smaller chromosomes.

Overall, our results indicate that the degree of genetic divergence in simulated data increases with the number of generation passes, and smaller chromosomes may be more susceptible to genetic drift.

#### Robustness - Variation in kinship estimates

The average interquartile ranges of N-REV rate estimates were calculated for chromosome one and 22 under both the WF (chr1: 0.024, chr22: 0.025) and Non-WF (chr1: 0.029, chr22: 0.021) models. Chromosome 1 had the widest range under the Non-WF model. However, the visual differences in box sizes were not noticeable in the accompanying Figure S10 and S11.

For chromosome 1, under the WF model, all N-REV rates ranged from 0.45 to 0.55 (Figure S10A). Most boxes showed no or one outlier, and only one box showed two outliers. On the other hand, under the Non-WF model, the N-REV rates ranged from 0.3 to 0.4 (Figure S10B) and most boxes (20 out of 27) showed no outlier, five showed one outlier, and two showed two outliers.

For chromosome 22, the N-REV rates under the WF model ranged from 0.24 to 0.35 (Figure S11A). Most boxes (23 out of 27) showed no outlier, followed by two boxes with one outlier, and two boxes with two outliers. Under the Non-WF model, the N-REV rates ranged from 0.18 to 0.25 (Figure S11B), with most boxes (17 out of 27) showing no outlier, six showing one outlier, and four showing two outliers.

Overall, these findings suggest that the estimates of N-REV rates varied slightly depending on the chromosome and the model used. However, the visual differences were not very pronounced, and most boxes had no outliers.

#### Ancestral accuracy - PCA visualization

Figure S8&S9 indicate that both chromosome one and 22 simulated data showed a similar trend of location shift in PC plots, regardless of the model used. Specifically, with 30 generation passes, the plots of simulated data almost perfectly overlapped with the referenced subpopulations when the input size was 90, with more than 95% of plots overlapping when the input size was 60, and the percentage reduced to 85%-90% when the input size was 30.

However, with 100 generation passes, the plots of simulated data deviated obviously towards the edge of the canvas, increasing the distance against other subpopulations. The location shift size increased by more than double with 300 generation passes, leading to a clear separation between the simulated and referenced subpopulations. The input size also played a role in modifying the speed of location shift inversely. Specifically, with 100 generation passes, an input size of 90 helped hold the position of most simulated data within the referenced area, but an input size of 30 led to half of the plots being separated from the referenced.

In summary of PCA plots, our simulation results demonstrate that the majority of ancestry can be maintained with a relatively small number of generation passes. However, after 300 generation passes, we observed a significant dispersion and location shift, leading to the simulated data plots becoming randomly spread out and out of control.

#### Ancestral accuracy - (M)ANOVA

The results of our (M)ANOVA analyses show that over 80% of P-values from F tests were smaller than 0.005, leading to rejections of the null hypothesis and indicating a significant separation between the simulated and referenced data (see Table S1&S2). MANOVA of top two PCs were the more powerful than ANOVA of one PC, although they were considered to be oversensitive. All MANOVA tests resulted in rejections for the simulated data with 100 or 300 generation passes. Under the WF model, only four YRI simulations with an input size of 60 and 30 generation passes survived MANOVA (Table S1). Under the Non-WF model, seven simulations in total, including four CHB simulations, two YRI simulations, and one GBR simulation, passed the test with an input size of 60 or 90 and 30 generation passes (Table S2). ANOVA tests of PC1 and PC2 alone were less sensitive and allowed more passes. Specifically, 32 WF simulations and 19 Non-WF simulations passed the ANOVA-PC1 test, and 34 WF simulations and 28 Non-WF simulations passed the ANOVA-PC2 test. Most of the ANOVA-PC1 test passes were from CHB and GBR simulations, and most of the ANOVA-PC2 test passes were from YRI simulations. Simulations with 30 generation passes had the highest passing rate of 0.21 and 0.15 under the WF and Non-WF models, respectively. Most simulations after 100 or 300 generation passes resulted in rejections. Simulations with an input size of 60 (WF: 0.13, Non-WF: 0.09) or 90 (WF: 0.09, Non-WF: 0.10) had similar passing rates, which were higher than that of simulations with an input size of 30 (WF: 0.05, Non-WF: 0.01). On average, simulation from WF model had a higher passing rate than Non-WF model (0.09 vs. 0.07). In general, our (M)ANOVA tests were found to be oversensitive in differentiating between similar ancestries. However, we observed that simulations with a larger input size and a smaller number of generation passes, particularly those using the WF model, were more likely to pass the test.

#### Relatedness - N-REV rate

Our analyses of N-REV rates consistently showed similar estimates across simulations with varying input sample sizes, numbers of generation passes, and reference ancestries. However, we observed that these rates differed when using different mating models and different chromosomes (see Figure S10&S11, panel A vs. B). Specifically, the average N-REV rates in simulations of chromosome one were 0.50 and 0.36 under the WF and Non-WF models, respectively, while in simulations of chromosome 22, they were 0.30 and 0.21, respectively. Simulations using the WF model and chromosome one had higher N-REV rates than those using the Non-WF model and chromosome 22.

### Inter-software practice

Following our extensive discussion and analyses (see Discussion-Best intro-software practice), we determined that simulations with an input size of 90 and 30 generation passes under the WF model were the best chromosome-level intro-software practice. These simulations consistently yielded similar ancestries and produced reliable estimates of N-REV rates. Therefore, we utilized above set of parameters for our exploration to the best chromosome-level intro-software practice.

#### PCA visualization

We assessed the performance of SLiM and HapGen approaches in maintaining PC position. We found that both approaches produced plots that overlapped with those of the referenced and anchor subpopulations, indicating that they were able to accurately capture the genetic variation present in these populations.

However, we also observed a slight location shift in the simulated plots generated by both approaches (see Figure S12). This shift was more noticeable in HapGen plots, which tended to be more concentrated than SLiM plots (Figure S12, panel A vs. B). Despite this shift, the simulated plots still maintained their overlap with the referenced and anchor subpopulations, suggesting that the location shift did not imply significant ancestry change.

We also noted that the dispersion of the SLiM plots was loose, causing the simulated plots to cover the plotting area of the referenced and anchor subpopulations when the output size exceeded 5,200. This dispersion may be attributed to the fact that SLiM is a forward-time simulation approach, which can result in a higher level of stochasticity in the simulations compared to HapGen, which is a resampling approach.

Overall, our findings suggest that both SLiM and HapGen approaches are suitable for simulating introgression events in human populations, with SLiM being more appropriate for larger output sizes and HapGen for smaller output sizes.

#### (M)ANOVA

Both simulation approaches, HapGen and SLiM, yielded significant P-values in oversensitive MANOVA tests, rejecting the null hypothesis of no separation between the simulated and referenced ancestries (Table S3). However, a few simulations passed the t-test in ANOVA of PC1 and PC2. With HapGen, one simulation with an output size of 500 passed the t-test in ANOVA of PC1, and another with an output size of 100 passed the t-test in ANOVA of PC2. With SLiM, four simulations with output sizes of 200, 500, 1300, and 30000 passed the t-test in ANOVA of PC1. All other simulations rejected the null hypothesis, suggesting significant differences in ancestry between the simulated and referenced populations.

Certainly, the (M)ANOVA tests were found to be oversensitive, as mentioned before. However, we were pleased to see that a few simulations using both HapGen and SLiM approaches still passed the ANOVA tests of one principal component (PC), indicating high level of similarity with the reference population.

#### Relatedness

Figure S13 showed that the N-REV rates from HapGen simulations were higher than those from SLiM when the output sizes were smaller than 13,000 (simulation ratio output/input = 144), and the opposite was observed when the output sizes were larger. The N-REV rates varied by output sizes in HapGen simulations, but they were consistent in SLiM simulations. Kinship coefficient estimates did not identify any REV in HapGen simulations when the output size was smaller or equal to 200. As the output size increased, the N-REV rate dropped rapidly. With the output sizes changing from 200 to 30,000 (200-fold), HapGen simulations’ N-REV rates decreased by 0.73. In contrast, SLiM simulations had little issue with N-REV rate drops. The increment of the output size from 100 to 30,000 (300-fold) only resulted in a 0.06 decrease in the N-REV rate.

These results suggest that the choice of simulation software and output size greatly impact the N-REV rate estimates. For small simulations, HapGen can still be considered the gold standard, but for larger simulations, SLiM may be a better option.

### Application

#### PCA

Results suggested that the size of the chromosome has an impact on the clustering in PCA plots, with smaller chromosomes resulting in looser clustering and larger standard deviations (See Figure S14). Whole-genome PCA showed the most concentrated plotting area, while chromosome 22’s PCA showed the loosest (Figure S14, panels A/B/C 1 vs. 4). Ancestral distances between different subpopulations were not obvious in identical PCA plots, but in normalized PCA (See Figure 5). The simulated GBR and CHB were closer to the referenced subpopulation than the IBS and JPT anchors, while the simulated YRI was farther from the referenced subpopulation than the EIN anchor, but not by much (Figure 5, panels A vs. D, B vs. E, C vs. F). Normalized PCA at the whole-genome level was more sensitive and showed a farther distance between tested and referenced data than chromosome-level analysis (Figure 5, plots pink vs. other colors).

**Figure 5:**
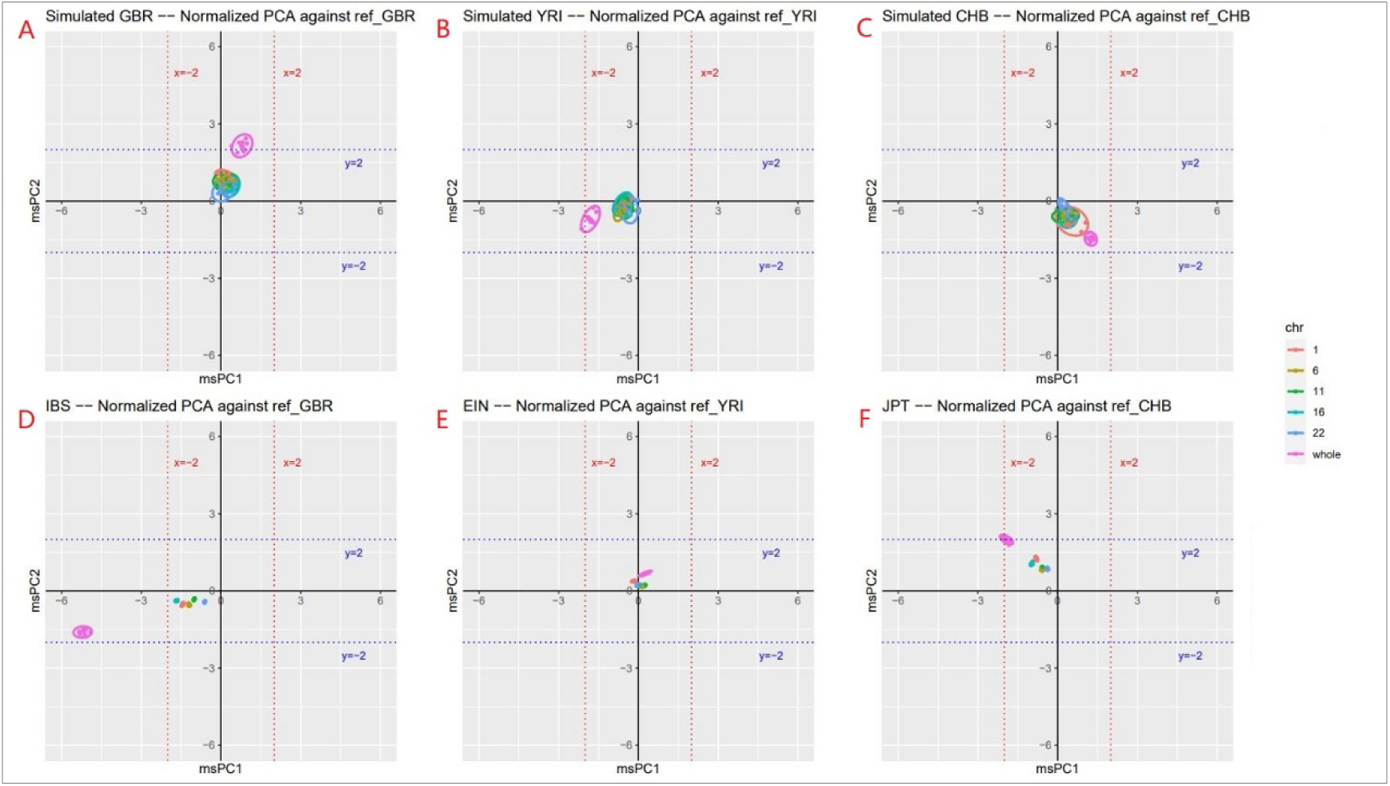
Plots of means of normalized PC1 & PC2 of simulated and anchor data. Plots summarized ten parallel simulations. **A**. Plots of simulated GBR. Normalization based on referenced GBR. **B**. Plots of simulated YRI. Normalization based on referenced YRI. **C**. Plots of simulated CHB. Normalization based on referenced CHB. **D**. Plots of IBS. Normalization based on referenced GBR. **E**. Plots of EIN. Normalization based on referenced YRI. **F**. Plots of JPT. Normalization based on referenced CHB. Dash lines mark 2x standard deviation from the mean PC1/PC2 of the referenced. Purple dots mark plots from whole-genome PCA. Other colored dots mark plots from chromosome-level PCA of 1, 6, 11, 16 and 22.

In (M)ANOVA results, tests using PCs derived from the whole genome were more sensitive than those using a single chromosome, and sensitivity varied by chromosome sizes and tested subpopulations (see Figure 6).

**Figure 6:**
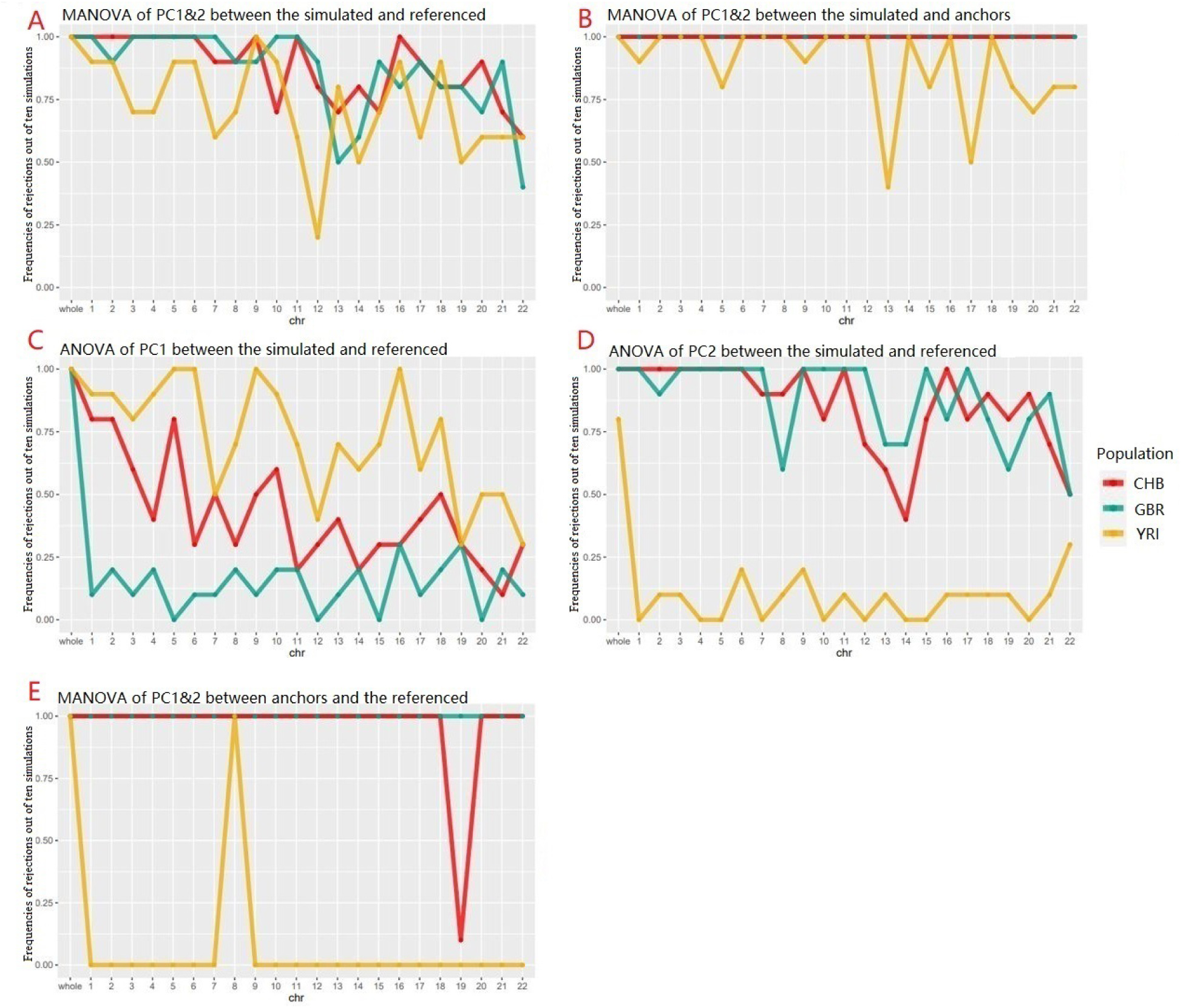
Frequencies of rejections to null hypotheses of PCA by chromosomes. Line charts represent frequency of rejections to the null hypotheses in F/t test of PCs in (M)ANOVA, across 10 parallel simulations. First column based on whole-genome PCA and others based on chromosome-level PCA. **A**. Comparison between the simulated and the referenced using MANOVA. **B**. Comparison between the simulated and anchors using MANOVA. **C**. Comparison between the simulated and the referenced using ANOVA of PC1. **D**. Comparison between the simulated and the referenced using ANOVA of PC2. **E**. Comparison between anchors and the referenced.

The results of (M)ANOVA tests that employed whole-genome PCs revealed significant differences between the simulated and referenced/anchor subpopulations across 10 parallel simulations, with only two exceptions (Figure 6, data point: leftmost). Specifically, ANOVA of PC2 failed to reject the null hypothesis in two out of ten simulations when comparing simulated YRI with the referenced YRI (Figure 6, panel D, data point: leftmost). Tests that used chromosome-level PCs rejected fewer null hypotheses, particularly for smaller chromosomes, resulting in lower concordance with tests that utilized whole-genome PCs ((Figure 6, data point: leftmost vs. others).

When comparing simulated and referenced subpopulations using chromosome-level MANOVA, the mean rejection rate for YRI (0.73) was lower than that of CHB (0.88) and GBR (0.87) (Figure 6A). This suggested that YRI was slightly easier to maintain the ancestral lineage during the simulation process.

When we compared the simulated subpopulations with the anchors, we observed that only a few chromosome-level MANOVA failed to reject the null in simulations of YRI, particularly when small chromosomes were used (Figure 6B). In contrast, all tests rejected the null in simulations of CHB and GBR. These results suggested that the ancestral differences between simulated CHB and JPT, and that between simulated GBR and IBS, were more significant than that between simulated YRI and EIN. Chromosome-level MANOVAs were found to be sensitive enough to identify the difference between simulated and anchor subpopulations.

Our analysis revealed that, on average, ANOVA of a single PC exhibited lower rejection rates compared to MANOVA of two PCs (Figure 6: C/D vs. A). Specifically, the rejection rates for ANOVA of PC1 and PC2 were 0.45 and 0.62, respectively, while the rejection rate for MANOVA of two PCs was 0.82. Interestingly, when we examined the concordance between the results obtained from ANOVA of PC1 and PC2, we found that GBR had the lowest rejection rates when using ANOVA of PC1 (0.17), while YRI had the lowest rejection rates when using ANOVA of PC2 (0.11). These findings suggest that ANOVA of a single PC may have limitations in accurately capturing the differences between subpopulations, particularly in populations with high levels of genetic diversity. Alternatively, combining multiple PCs in MANOVA may provide a more comprehensive understanding of the differences between subpopulations.

We performed MANOVA between the referenced and anchor subpopulations (Figure 6E) and found that only MANOVA using chromosome 8 exhibited the same rejection rate as using the whole genome to differentiate the referenced YRI from EIN. Moreover, only chromosome 19 yielded different results (concordance = 0.1) from using the whole genome to identify the referenced CHB from JPT. Upon further examination, we concluded that the genetic differences between the referenced YRI and EIN were primarily distributed on chromosome 8. Similarly, chromosome 19 also had a minor contribution to the genetic differences between the referenced CHB and JPT, while the primary differences were found across other chromosomes. These results suggest that certain chromosomes may be more informative than others in distinguishing between subpopulations and that a comprehensive analysis of multiple chromosomes may yield a more accurate assessment of genetic differences between subpopulations.

Moreover, our analysis indicated that the (M)ANOVA results obtained from chromosome one were highly consistent with those obtained from the whole genome analysis in general cases with no selection effect (Figure 6, panels A and B). This suggests that chromosome one may contain a significant amount of genetic variation that is representative of the entire genome and that it may be particularly informative in studying genetic differences between subpopulations. However, it is important to note that in some special cases, where tests were not powerful enough or genetic differences between two subpopulations were not evenly distributed across all chromosomes, whole-genome tests were irreplaceable by chromosome-level tests. For instance, our analysis showed that MANOVA using chromosome one was unable to identify the genetic differences between YRI and EIN. In such cases, a PCA of the entire genome may be necessary to accurately identify genetic differences between subpopulations.

#### N-REV rate

Our analysis showed that whole-genome N-REV rate estimates were significantly more accurate than chromosome-level estimates (see Figure 7, leftmost vs. others). Only whole-genome estimates fell within the confidence interval of observed true values, whereas chromosome-level estimates were all biased and consistently lower than true values. Moreover, as the chromosome size decreased, the N-REV rate estimates obtained from chromosome-level analysis became increasingly overaggressive in identifying REVs, suggesting a loss of precision in estimation. Notably, the N-REV rate estimates were consistent across the three subpopulations tested. Taken together, our findings indicate that chromosome-level kinship estimates should be avoided due to their inherent bias and reduced accuracy compared to whole-genome estimates.

**Figure 7:**
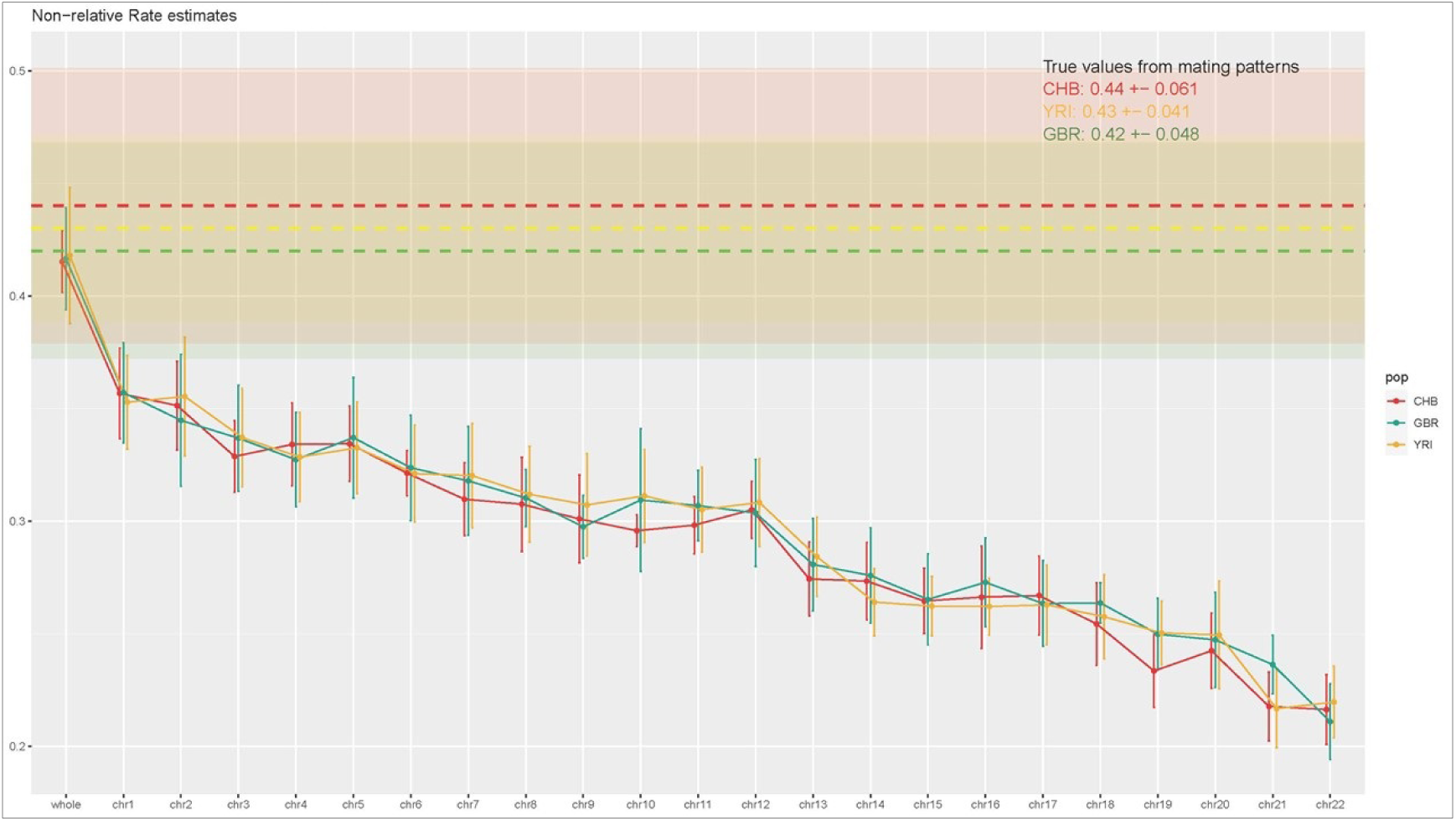
N-REV rate estimates by chromosomes. Dash lines are mean N-REV rates derived from mating patterns (true values) across 10 parallel simulations and shadows are error bars. Solid lines are estimated means and error bars based on kinship analyses. First column present statistics derived from whole-genome analyses and others are derived from chromosome-level analyses. CHB colored in red. GBR colored in green. YRI colored in yellow.

## Discussion

The primary aim of this study was to introduce a novel and standardized method for generating pseudo-genotype data via a random mating system using SLiM. Our approach for chromosome-level simulation was found to be comparable to the widely-used HapGen method. Specifically, our approach using SLiM proved to be more effective in simulating large sample sizes, while HapGen remains the gold-standard for simulating smaller samples.

In addition to this primary objective, we also evaluated the sensitivity of chromosome-level quality control (QC) statistics using our whole-genome simulation approach. Our application of this approach allowed us to assess the effect modification of population structure by chromosomes, which has not been successfully demonstrated in previous whole-genome simulations. Our results indicated that chromosome-level PCA and kinship estimates were less reliable than whole-genome statistics and as population structure differs along the genome, which aligns with previous reports (6).

### Summary of findings

This study presents several important findings related to the simulation of pseudo-genotype data using the SLiM program. The results indicate that increasing the input size of the simulation improves robustness, ancestral accuracy, and relatedness, with marginal utility beyond an input size of 60. Simulating more than 30 generations does not help relatedness but creates an ancestral shift and less robustness. The study recommends using the WF rather than Non-WF model for chromosome-level simulation in cases of lower relatedness, and an input size of 60 or greater with 30 generation passes for simulations using SLiM.

In terms of simulation approaches, the study found that both SLiM and HapGen were capable of simulating samples up to 333x the input size while maintaining the same ancestry as the referenced populations. SLiM outperformed HapGen regarding relatedness when the simulation ratio is high, while HapGen is still a gold standard if the application only needs a small to mediocre sample size.

The study also found that chromosome-level PCA was less sensitive than whole-genome PCA in differentiating ancestries, and sensitivity decreased with a decrement in chromosome size. Similarly, chromosome-level kinship analysis induced bias of N-REV estimates and was more aggressive in identifying REVs with a decrement in chromosome size. We verified differences in population structure across the genome by observing certain chromosomes, such as chromosome 8 and 19, differently informative than others in distinguishing specific subpopulations.

Overall, the study provides valuable insights into the simulation of pseudo-genotype data using the SLiM program and makes recommendations regarding simulation approaches and input parameters. The findings highlight the importance of considering the target output size and research interest when choosing a simulation tool. Last but not least, we provices supports to caution in using chromosome-level PCA and kinship estimates.

### Strengths

We are the first to date to provide a standardized whole-genome genotype simulation approach. Previous approaches limited to regional or chromosome-level simulation only. Our approach yielded the same ancestral accuracy of simulation as the current gold standard. Different from HapGen, our approach used a mating system and held a consistent N-REV rate even simulating tens of thousands of genotypes. Additionally, HapGen stopped updating long time ago, but SLiM is still under maintenance and updating and we will update our approach accordingly.

In our application, the identification of certain chromosomes as differently informative in distinguishing specific subpopulations is a valuable finding that can aid future research in this area. Ultimately, our cautionary recommendation regarding the use of chromosome-level PCA and kinship estimates is important for ensuring the accuracy of such analyses.

### Limitations

The first limitation of the study is the high number of REVs produced due to the inbreeding in the mating system, limit the use of our simulation for small sample sizes. The second limitation is the longer running time compared to existing tools like HapGen, as our approach take more time modeling the complicated mating system. The third limitation is the inability of the approach to accurately simulate extremely rare variants, as they are likely to be lost due to allele frequency fluctuations over generations of mating.

### Future works

#### Advanced simulation

The present study provided a simplified simulation approach with unisexual mating, uniform recombination, and neutral mutation. However, there is significant potential for improving the simulation’s accuracy by incorporating more customized settings. For example, the use of hotspots in the human genome to model complex recombination and mutation would better maintain the genetic structure during simulation. Additionally, the latest version of SLiM includes a sexual mating system that should be evaluated in future work.

It is important to note that all mating simulations performed in this study utilized reference data within a specific subpopulation. Therefore, it would be interesting to compare the quality of results obtained from an admixed simulation to those we have obtained. This would allow us to gain a better understanding of any differences in the genetic structure and how this may affect future analyses.

#### More applications

Our novel simulation approach has the potential to be widely adopted for validating new statistical methods. Developers can apply this approach to generate tens of thousands of pseudo-genotype data using the 1000 Genomes Project. For our specific research interest, we plan to utilize this approach to evaluate a new statistic for genetic imputation. By doing so, we can assess the effectiveness of the method in a controlled environment and better understand its performance under various conditions. This approach can also be extended to evaluate other statistical methods and their potential applications.

## Materials and methods

### Data and formats

The initial data input used in this study was the 1000-genome project (1KGP; URL: https://www.internationalgenome.org/category/ftp/), a publicly available, whole-genome sequencing dataset of diverse populations (7). A subset of the 1KGP was selected by performing a PCA (Figure S1), and individuals from Han Chinese in Beijing, China (CHB, N=103), British from England and Scotland (GBR, N=91), and Yoruba in Ibadan, Nigeria (YRI, N=108) were chosen based on the PCA as distinct ancestral subpopulations capturing the three primary continental clusters. These subpopulations were chosen as they represent most GWAS/WGS samples and continental ancestry (8).

The simulation and application process utilized three data formats: FASTQ, variant call format (VCF), and Plink binaries. The 1000-genome data is publicly available in the FASTQ and VCF encoding formats. FASTQ is a text-based format used to store biological sequences and the corresponding quality scores (9). The sequence letter and quality score are encoded with a single ASCII character for brevity. The VCF format specifies the form of a text file used in bioinformatics for storing gene sequence variations (4). Variants were stored along with a reference genome using VCF. Plink binary files include a binary genotype file, a genetic map, and a family pedigree file (.bed, .bim, .fam) (10). These binaries record the same information as VCFs but are more compressed to reduce data storage.

To create the whole-genome-level pseudo genotype data, SLiM was used, taking VCF and FASTQ as input nucleotide data and outputting the VCF format. For storage efficiency, the simulated data was converted to Plink binary formats for QC and downstream analyses.

### Tools

Within the study, the following tools were used: Plink(v1.9/v2.0), R(v4.1.2), SLiM(v3.3), VCFtools(v0.1.13), SAMtools(v1.13), EIGENSOFT(v8.0.0), and KING(v2.2.7) (4,5,10–15). During the study, we employed VCFtools and SAMtools to manage and clean the sequencing data. We utilized only the basic features of Plink v2.0, and primary quality control (QC) was executed using Plink v1.9. SLiM v3.3 was utilized throughout the entire simulation process. For the QC and analysis of output data, EIGENSOFT was used for both exact and approximate principal component analysis, while KING was utilized to identify related individuals through kinship analysis.

### Code resource

We shared step-by-step suggestions for customized simulations, quality control steps, and evaluations. All codes can be found on our GitHub: (https://github.com/zxc307/GWAS_simulation_handbook).

### Reference QC

In this publication, we presented a reference QC checklist for the scenarios we addressed, including: 1. genome reference build, 2. call rate, 3. minor allele frequency (MAF), 4. INDELs, 5. multi-allelic markers, 6. duplicate loci, 7. hardy-Weinberg, 8. ancestry, 9. independence, 10. non-nucleotide loci (e.g., unknown/aMino/puRine). We utilized genome consortium human builds 37, and downloaded the pre-cleaned loci data from the 1KGP FTP server, excluding cites with call rates lower than 95% and all duplicates. The input data passed Plink’s default hardy-Weinberg test (P <= 0.01), and we performed the QC checklist for the remaining conditions 3-5, 9-10. We excluded all monomorphic, INDELs, multi-allelic, and NMR variants, as well as any relatives described in the profile of 1KGP (16).

### Simulation with SliM

SliM provided us with a powerful set of features and algorithms to simulate our data. Below we outline the key settings, features, and algorithms used in our study.

#### Mutation and recombination

In reality, ancestral populations, gender, and genomic regions all contribute to differences in the mutation and recombination rates of meiosis (17,18). These rates also influence selection in nature. While SliM allows for modeling this complex distribution of mutation and recombination, it was not relevant to our objectives to induce more selection effects. Therefore, for simplicity, we assumed that all recombination and mutation events during the mating process were uniformly distributed across all polymorphic variants in the whole genome, thus minimizing the selection effect. We set the mutation effect to be neutral with a rate of 1×10^−7^, and the recombination rate was set to 1×10^−8^ per generation. As a result of the assumption, our set of mutation and recombination events was sensitive to the distribution of variants across the autosomal genome, with higher frequencies of mutations and recombination events in SNP-dense regions compared to SNP deserts.

#### Mating system

In our simulation, we used a forward mating system in which chromosomes were randomly distributed during pseudo meiosis. The simulation included only autosomal chromosomes, and mating was unisexual and biparental. All input data were marked as the first generation, and each mating event involved the selection of two founders from the pool, with no regard to sex. Recombination and mutation were processed on the sampled founders, and an offspring was generated. A visual example of a one-generation pass is provided in Figure 1. The simulation excluded all monomorphic sites, and two haploids recombined at randomly simulated sites, with one copy sampled thereafter. Mutations were then added to the copy, and the haploid was paired with another prepared one to form an offspring. In order to mitigate the negative impacts of inbreeding, we implemented a strategy whereby the maximum number of matings at each generation was limited to the size of the available pool of samples. All matings were processed independently at individual level.

#### Mating pattern

For the chromosome-level simulation, we implemented randomized mating patterns for simulations on individual chromosomes and in parallel runs. We kept track of every mating pattern, including the founder pairs, offspring, and sampling indicators. In each run of the whole-genome simulation, we applied the same pattern to all chromosomes to ensure that mating pairs remained consistent throughout the genome, thereby preserving cross-chromosome associations.

#### WF model

According to the Wright-Fisher model, the population simulation consists of three stages. In the first stage, the population undergoes a boosting phase. At this stage, all founders and offspring are kept, resulting in a doubling of the sample size after each generation pass. Once the sample size exceeds the desired output size, the simulation proceeds to the conjunction stage. Here, random sampling is performed to obtain a sample size that matches the specified output size. After this, the simulation moves to the population maintenance stage, which is the third and final stage. In this stage, the number of mating events is equal to the output sample size, and only offspring are retained while parental samples are excluded.

#### Non-WF model

The Non-Wright-Fisher (Non-WF) model is a dynamic ecology that comprises three stages, with the first two stages being similar to those in the Wright-Fisher (WF) model. However, the third stage in the Non-WF model differs from that of the WF model. We have highlighted the key differences between the third stages of the two models in Figure 2. In the third stage of the Non-WF model, the mating number equals the sample size, and we combine the offspring with the older generation and filter them using SLiM’s default binary fitness model at each generation pass. The fitness model’s details can be found in section 8 of the SLiM manual (19). As we filter in every generation at this stage, older samples undergo more filtrations, resulting in an age-sensitive model of cumulative fitness, where younger generations are more likely to survive. Not only do the procedures of the third stage differ in the two models, but the SLiM modules also differ technically. The WF module, which was developed in 2013, did not support the recording of mating patterns for the whole-genome simulation. However, the Non-WF module, which was released in 2019 and subsequently updated, was more customizable and covered new features, including pattern recording that enabled whole-genome simulation.

### QC statistics

The two most commonly used quality control (QC) statistics in genome-wide association studies (GWAS) are principal component analysis (PCA) and kinship estimates. We utilized these assessments as key measures throughout our study and provide several details about them below.

#### Reference array

To create a quality control (QC) reference array, we pruned the 1KGP phase 3 data using a sliding window of 150 variants and a shift of 15 variants between windows, with an independent pairwise r^2^ threshold of 0.2. We also excluded uncommon variants with a minor allele frequency (MAF) less than 0.05. We applied this reference array to our PCAs and kinship analyses in the study.

#### PCA - Hypothesis test

We employed PCA to assess whether the ancestry of our tested sample matched that of the reference samples. Our null hypothesis was that the tested and reference samples shared the same ancestry and, therefore, the same distribution of principal components (PCs). We evaluated this hypothesis through visualization of identical PCA plots, normalized PCA plots, and (M)ANOVA tests.

#### PCA - Anchors

To improve visualization of the distance between our tested and reference subpopulations, we utilized anchors. We selected Esan in Nigeria (EIN), Japanese in Tokyo (JPT), and Iberian Populations in Spain (IBS) as our anchors, as they are the closest subpopulations to the reference samples of Yoruba in Ibadan, Nigeria (YRI), Han Chinese in Beijing, China (CHB), and British in England and Scotland (GBR) from the 1KGP dataset.

#### PCA - Normalized PCA

Based on our analysis of histograms of the top two PCs of our reference subpopulations (Figure S2), we made the assumption that these PCs followed normal distributions. Therefore, under the null hypothesis, tested samples should have fallen within the normal distribution of the reference samples. In addition to evaluating identical PCs, we also performed normalization of the tested data’s PCs, as

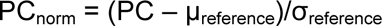

where µ_reference_ was the mean of PC_reference_ and σ_reference_ was the standard deviation of PC_reference_. Our expectation was that plots of normalized PCs generated from the test data would display the distance between the tested and reference samples in the same scale.

#### PCA - (M)ANOVA tests

To test our hypothesis, we employed multivariate analysis of variance (MANOVA) for the top two PCs and analysis of variance (ANOVA) for each PC separately (20). Since we conducted multiple tests across ten parallel simulations, we used Bonferroni correction to adjust the type I error rate to 0.005 for the F statistical tests of (M)ANOVA (21). A p-value less than 0.005 would result in rejecting the null hypothesis, indicating that the tested samples have a different ancestry than the reference samples.

#### Relatedness - True families from mating records

We utilized mating records to calculate the true values of relatedness, and presented a family tree (Figure S3) and clusters of related samples (Figure S4) from one simulation run. This example consisted of 30 families ranging in size from two to 25 individuals. To rescue unrelated samples from these families, we employed two approaches. In the conservative approach, we rescued one individual from each family, while in the aggressive approach, we rescued multiple individuals from large families (greater than three) and as many non-second-degree or closer relatives (N-REVs) as possible. In the example, we were able to rescue 30 and 52 individuals using the conservative and aggressive approaches, respectively. To determine the final number of rescued individuals for downstream analysis, we took an average of these numbers, resulting in 46 individuals for this example. The same calculations were applied to all simulations.

#### Relatedness - Kinship estimate

To estimate kinship coefficients, we utilized KING’s fast and integrated relationship inference method, using identical by state to predict identical by descent and calculate the kinship coefficient. We employed a cutoff of 0.0884 to identify second-degree or closer relatives (REVs). For pairs of samples with kinship coefficients greater than the threshold, we excluded one member using kinship coefficient matrices. After the kinship quality control (QC) process, we considered all surviving samples as N-REVs based on our estimates.

### Best practice

We conducted a chromosome-level evaluation of Non-WF, WF, and HapGen simulations, as only the Non-WF model technically supports whole-genome simulation. Initially, we examined the optimal practices within SliM by varying parameters. Subsequently, we compared the performance of SliM with that of HapGen.

#### Intro-software practice

Our study conducted a sensitivity analysis of simulation quality within SLiM by varying several simulation parameters to simulate a fixed number of individuals (200). These parameters included the simulation model setting (WF and Non-WF), ancestry of the input reference panel (CHB, GBR, and YRI), number of generation passes (30, 100, and 300), and input sample size (30, 60, and 90).

We evaluated simulation quality in terms of robustness, ancestral accuracy, and percentage of N-REV pass. We presumed that the validity of chromosome-level QC statistics positively correlated with the chromosome size, with the smallest chromosome (chromosome 22) having the largest potential for variation in simulated statistics with the smallest input sample size of 30, longest generation passes of 300, and under the WF model.

To evaluate robustness, we replicated all simulations of chromosome 22 ten times with different random seeds and summarized the standard deviations of key statistics. To assess ancestral accuracy, we used PCs from chromosome one simulated data, as they were presumed to be more valid than PCs from smaller chromosomes. We visually assessed the accuracy using PC plots and conducted (M)ANOVA of the top two PCs. The percentage of N-REV pass reflects the simulation efficiency, as REV samples need to be excluded after simulation in some cases. We calculated the N-REV rate based on kinship coefficient estimates.

#### Inter-software practice

We conducted parameter optimization for SLiM based on best practices in the field and then compared its performance with that of HapGen. Specifically, we simulated chromosome 15 data for various sample sizes (ranging from 100 to 30,000 individuals) in the GBR subpopulation using both approaches. We assessed simulation quality in terms of ancestral accuracy and relativeness, and compared data from the two methods. We used the same assessment criteria as in the intro-software practice, including PCA and N-REV rate calculations.

### Application

As of December 2022, SLiM is currently the only software that enables whole-genome simulation while retaining cross-chromosome associations. Thus, we were unable to compare our simulation results with those of another method for evaluation purposes. Instead, we demonstrated the practical application of our simulation results in downstream analyses.

#### Evaluation of chromosome-level QC statistics

Recent studies have suggested that the impact of population structure can vary across the genome (6). While commonly used quality control (QC) measures like PCs and kinship coefficients have been developed and validated for whole-genome data, their validity when applied to a single chromosome is uncertain. To address this, we conducted whole-genome simulations to evaluate the sensitivity of chromosome-level QC statistics across the genome.

We employed a Non-WF model to simulate genetic data and set the input population size to 60 with an output size of 200 individuals. We performed ten parallel simulations with 30 generations using three different subpopulation references, including CHB, GBR, and YRI.

To evaluate the quality of simulations at both the whole-genome and chromosome levels, we utilized the same quality assessment methods from our best practice exploration process. Then, we compared chromosome-level principal component (PC) and kinship coefficient estimates to their whole-genome-level counterparts and true values, if applicable.

## Acknowledgments

I would like to take this opportunity to express my sincere gratitude to all those who have contributed to the publication of my work. Firstly, I would like to extend my heartfelt thanks to my supervisor, Dr. Schumacher, for his invaluable guidance, support, and feedback throughout the entire project. Without his mentorship, this publication would not have been possible. I would also like to express my deep appreciation to the developers of SliM, KING, and Plink2, Dr. Benjamin C. Haller at Cornell University, Dr. Wei-Min Chen at the University of Virginia, and Dr. Christopher Chang at GRAIL, Inc. Their software has been critical to the success of my project, and their timely responses to my queries have been incredibly helpful. Their insightful comments and suggestions have helped me to refine my approach and bring this project to completion.

## SUPPLEMENTAL MATERIALS

**Supplementary Table 1:** Summary of (M)ANOVA tests in intro-software practice under WF model

| MANOVA\|ANOVA-PC1\|ANOVA-PC2 |  |  |  |  |
| --- | --- | --- | --- | --- |
| Sub-population=CHB |  |  |  |  |
|  |  | n_generation |  |  |
|  |  | 30 | 100 | 300 |
|  | 30 | 10\|10\|10 | 10\|10\|10 | 10\|10\|10 |
| N_input | 60 | 10\|1\|10 | 10\|10\|10 | 10\|10\|10 |
|  | 90 | 10\|2\|10 | 10\|8\|10 | 10\|10\|10 |
| Sub-population=YRI |  |  |  |  |
|  |  | n_generation |  |  |
|  |  | 30 | 100 | 300 |
|  | 30 | 10\|10\|3 | 10\|10\|8 | 10\|10\|10 |
| N_input | 60 | 6\|7\|1 | 10\|10\|6 | 10\|10\|9 |
|  | 90 | 10\|10\|1 | 10\|10\|8 | 10\|10\|10 |
| Sub-population=GBR |  |  |  |  |
|  |  | n_generation |  |  |
|  |  | 30 | 100 | 300 |
|  | 30 | 10\|6\|10 | 10\|10\|10 | 10\|10\|10 |
| N_input | 60 | 10\|7\|10 | 10\|9\|10 | 10\|10\|10 |
|  | 90 | 10\|8\|10 | 10\|10\|10 | 10\|10\|10 |

Note: Each cell in the table represents count of rejections to null hypothesis of PCA tests using chromosome one data and (M)ANOVA tests, across 10 parallel simulations.

**Supplementary Table 2:** Summary of (M)ANOVA tests in intro-software practice under Non-WF model

| MANOVA\|ANOVA-PC1\|ANOVA-PC2 |  |  |  |  |
| --- | --- | --- | --- | --- |
| Sub-population=CHB |  |  |  |  |
|  |  | n_generation |  |  |
|  |  | 30 | 100 | 300 |
|  | 30 | 10\|10\|10 | 10\|10\|10 | 10\|10\|10 |
| N_input | 60 | 8\|7\|8 | 10\|10\|9 | 10\|9\|9 |
|  | 90 | 8\|7\|6 | 10\|8\|9 | 10\|10\|10 |
| Sub-population=YRI |  |  |  |  |
|  |  | n_generation |  |  |
|  |  | 30 | 100 | 300 |
|  | 30 | 10\|10\|6 | 10\|10\|10 | 10\|10\|10 |
| N_input | 60 | 10\|10\|5 | 10\|10\|9 | 10\|10\|10 |
|  | 90 | 8\|8\|4 | 10\|10\|8 | 10\|10\|10 |
| Sub-population=GBR |  |  |  |  |
|  |  | n_generation |  |  |
|  |  | 30 | 100 | 300 |
|  | 30 | 10\|10\|10 | 10\|10\|10 | 10\|10\|10 |
| N_input | 60 | 9\|7\|9 | 10\|10\|10 | 10\|8\|10 |
|  | 90 | 10\|9\|10 | 10\|10\|10 | 10\|8\|10 |

Note: Each cell in the table represents count of rejections to null hypothesis of PCA tests using chromosome one data and (M)ANOVA tests, across 10 parallel simulations.

**Supplementary Table 3:**
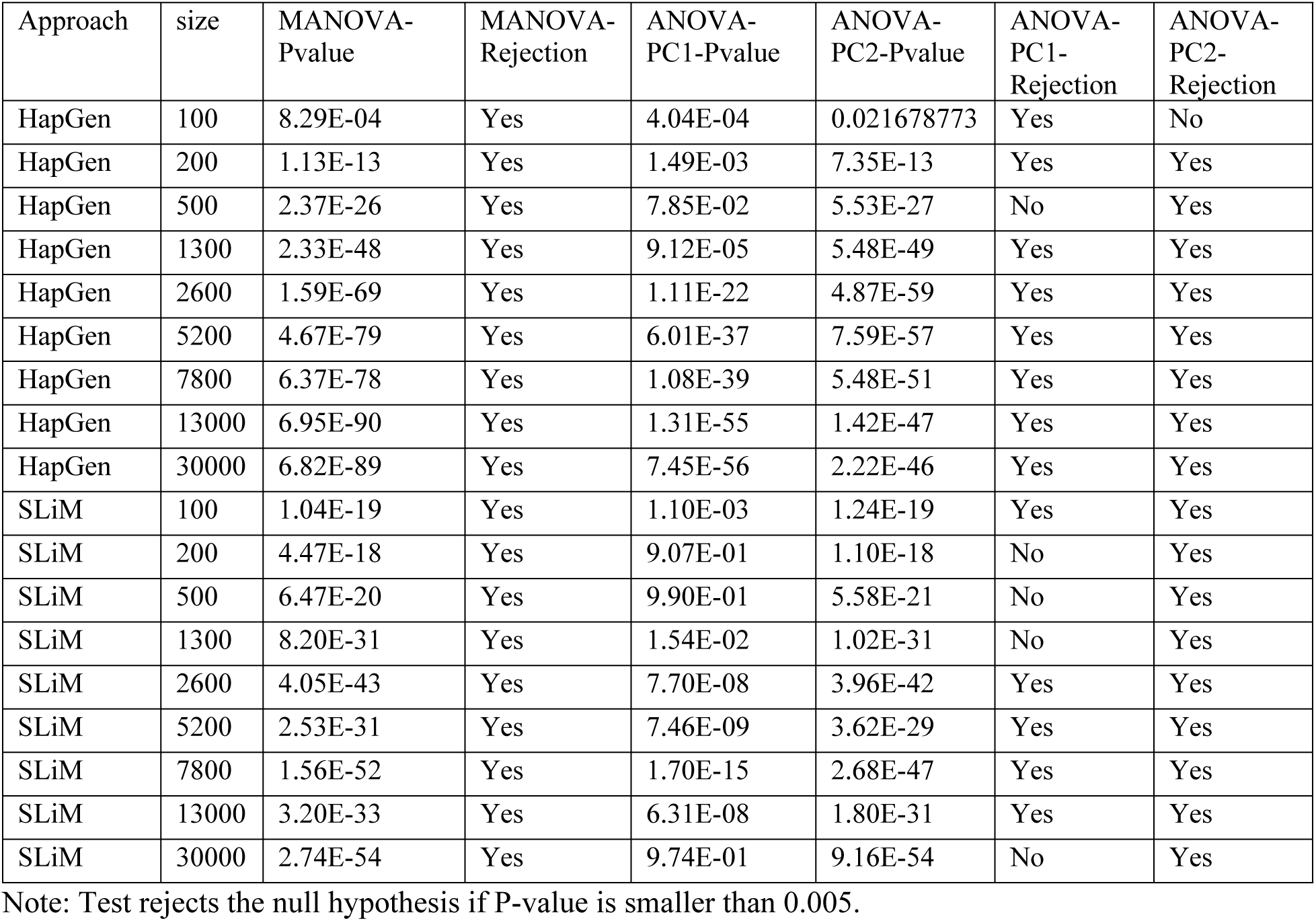
Summary of (M)ANOVA of inter-software practice

| Approach | size | MANOVA-Pvalue | MANOVA-Rejection | ANOVA-PC1-Pvalue | ANOVA-PC2-Pvalue | ANOVA-PC1-Rejection | ANOVA-PC2-Rejection |
| --- | --- | --- | --- | --- | --- | --- | --- |
| HapGen | 100 | 8.29E-04 | Yes | 4.04E-04 | 0.021678773 | Yes | No |
| HapGen | 200 | 1.13E-13 | Yes | 1.49E-03 | 7.35E-13 | Yes | Yes |
| HapGen | 500 | 2.37E-26 | Yes | 7.85E-02 | 5.53E-27 | No | Yes |
| HapGen | 1300 | 2.33E-48 | Yes | 9.12E-05 | 5.48E-49 | Yes | Yes |
| HapGen | 2600 | 1.59E-69 | Yes | 1.11E-22 | 4.87E-59 | Yes | Yes |
| HapGen | 5200 | 4.67E-79 | Yes | 6.01E-37 | 7.59E-57 | Yes | Yes |
| HapGen | 7800 | 6.37E-78 | Yes | 1.08E-39 | 5.48E-51 | Yes | Yes |
| HapGen | 13000 | 6.95E-90 | Yes | 1.31E-55 | 1.42E-47 | Yes | Yes |
| HapGen | 30000 | 6.82E-89 | Yes | 7.45E-56 | 2.22E-46 | Yes | Yes |
| SLiM | 100 | 1.04E-19 | Yes | 1.10E-03 | 1.24E-19 | Yes | Yes |
| SLiM | 200 | 4.47E-18 | Yes | 9.07E-01 | 1.10E-18 | No | Yes |
| SLiM | 500 | 6.47E-20 | Yes | 9.90E-01 | 5.58E-21 | No | Yes |
| SLiM | 1300 | 8.20E-31 | Yes | 1.54E-02 | 1.02E-31 | No | Yes |
| SLiM | 2600 | 4.05E-43 | Yes | 7.70E-08 | 3.96E-42 | Yes | Yes |
| SLiM | 5200 | 2.53E-31 | Yes | 7.46E-09 | 3.62E-29 | Yes | Yes |
| SLiM | 7800 | 1.56E-52 | Yes | 1.70E-15 | 2.68E-47 | Yes | Yes |
| SLiM | 13000 | 3.20E-33 | Yes | 6.31E-08 | 1.80E-31 | Yes | Yes |
| SLiM | 30000 | 2.74E-54 | Yes | 9.74E-01 | 9.16E-54 | No | Yes |

Note: Test rejects the null hypothesis if P-value is smaller than 0.005.

**Supplementary Figure 1:**
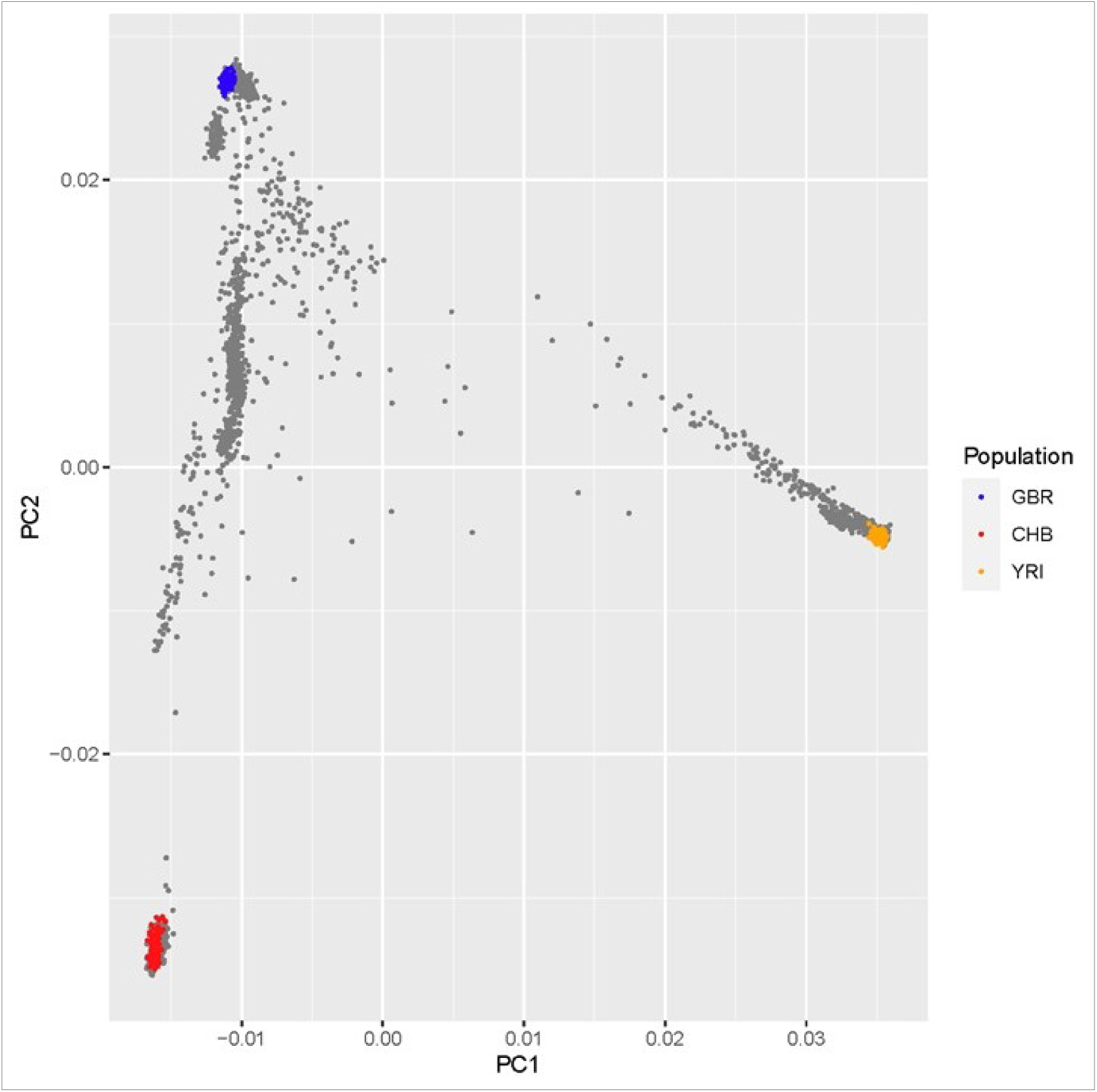
Plots of PC1 vs PC2 of 1000-Genome data (n=2,504) GBR: British from England and Scotland, in blue. CHB: Han Chinese in Beijing, China, in red. YRI: Yoruba in Ibadan, Nigeria, in orange. Other populations are in grey.

**Supplementary Figure 2:**
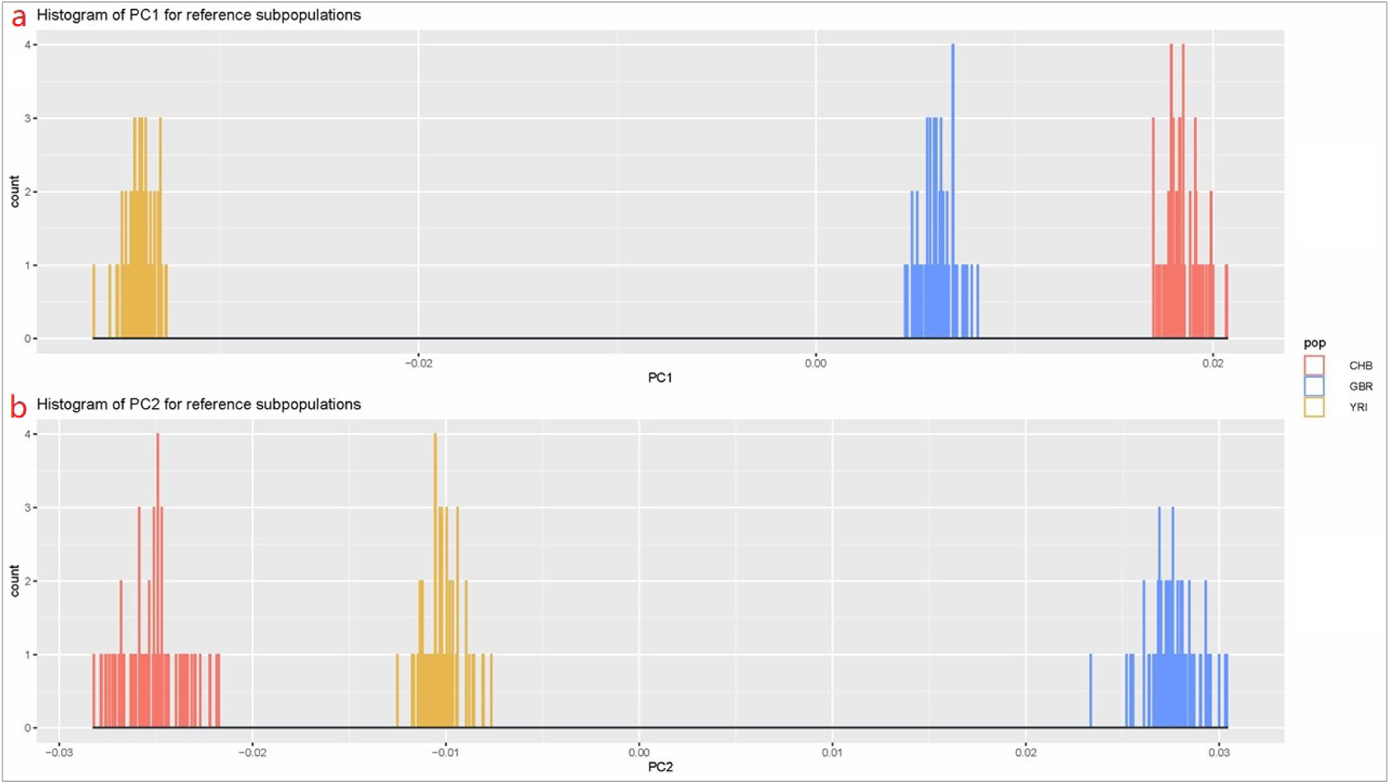
Histograms of first two PCs for reference subpopulations. **a**. Histogram of PC1. **b**. Histogram of PC2. CHB: CHB subpopulation reference in 1KGP, in red. GBR: GBR subpopulation reference in 1KGP, in green. YRI: YRI subpopulation reference in 1KGP, in blue.

**Supplementary Figure 3:**
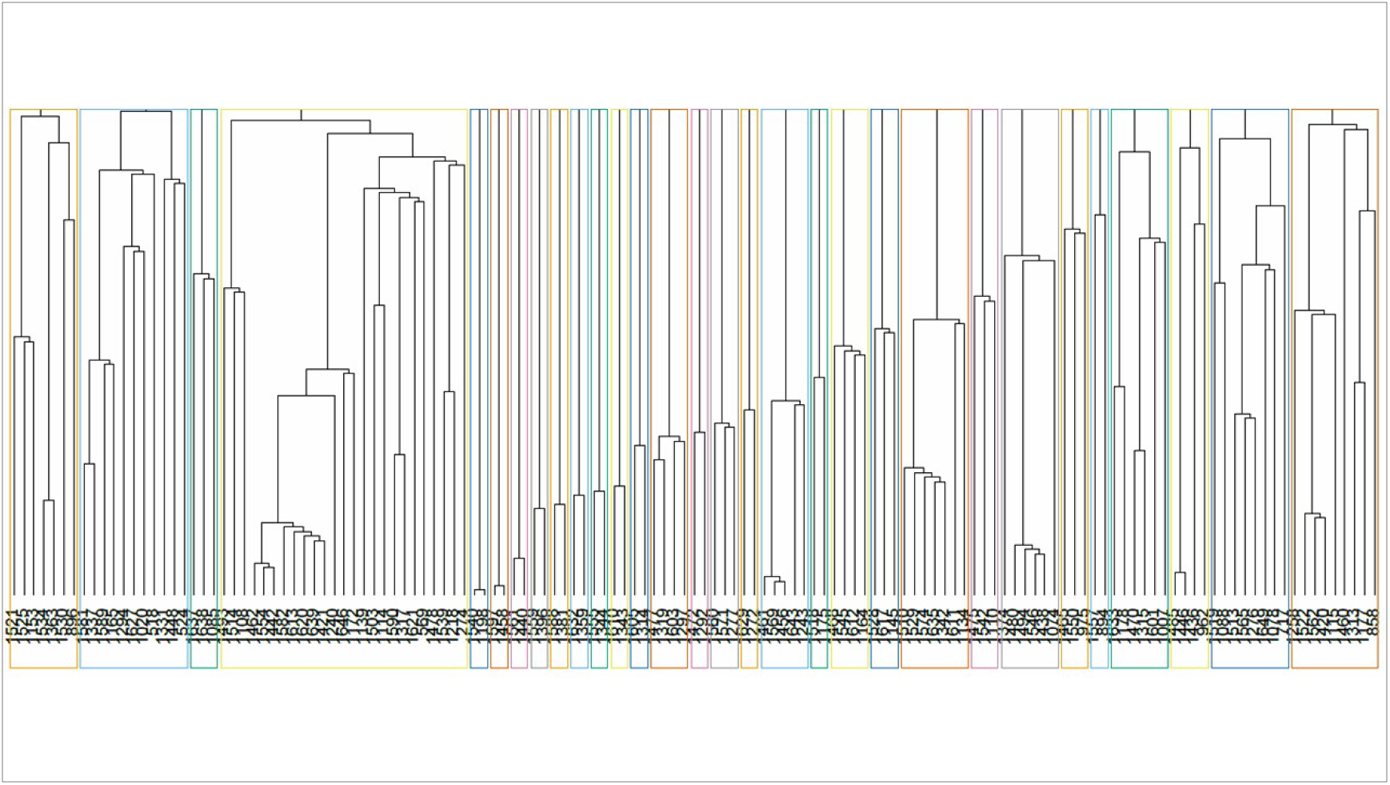
Family tree. An example from #1 simulation using 30 GBR as the reference after mating of 30 generations. Each number on the bottom represents one individual. Each colored rectangle represents one family cluster. Every branch represents a generation pass.

**Supplementary Figure 4:**
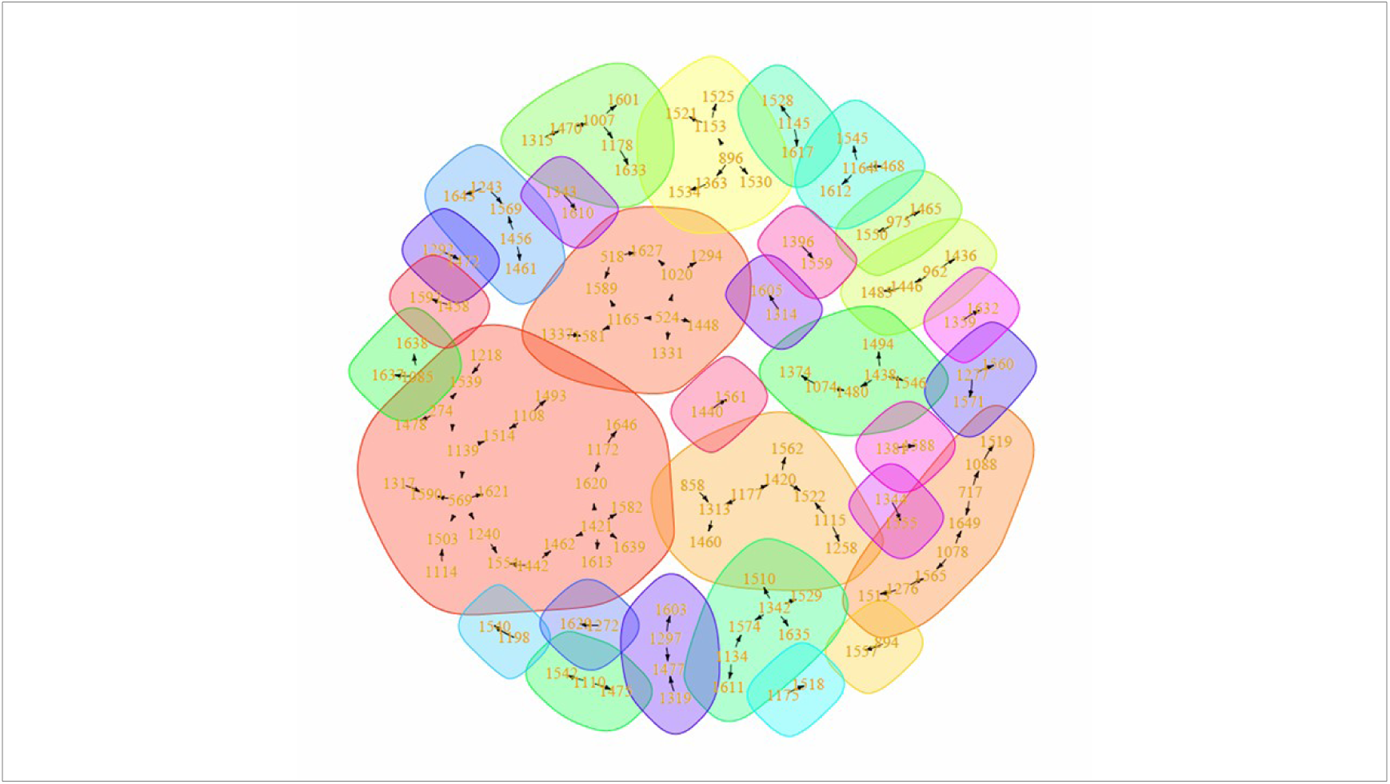
Family cluster. An example from #1 simulation using 30 GBR as the reference after mating of 30 generations. Each number represents one individual. Colored shadows represent families. Each arrow represents a generation pass from a founder to offspring

**Supplementary Figure 5:**
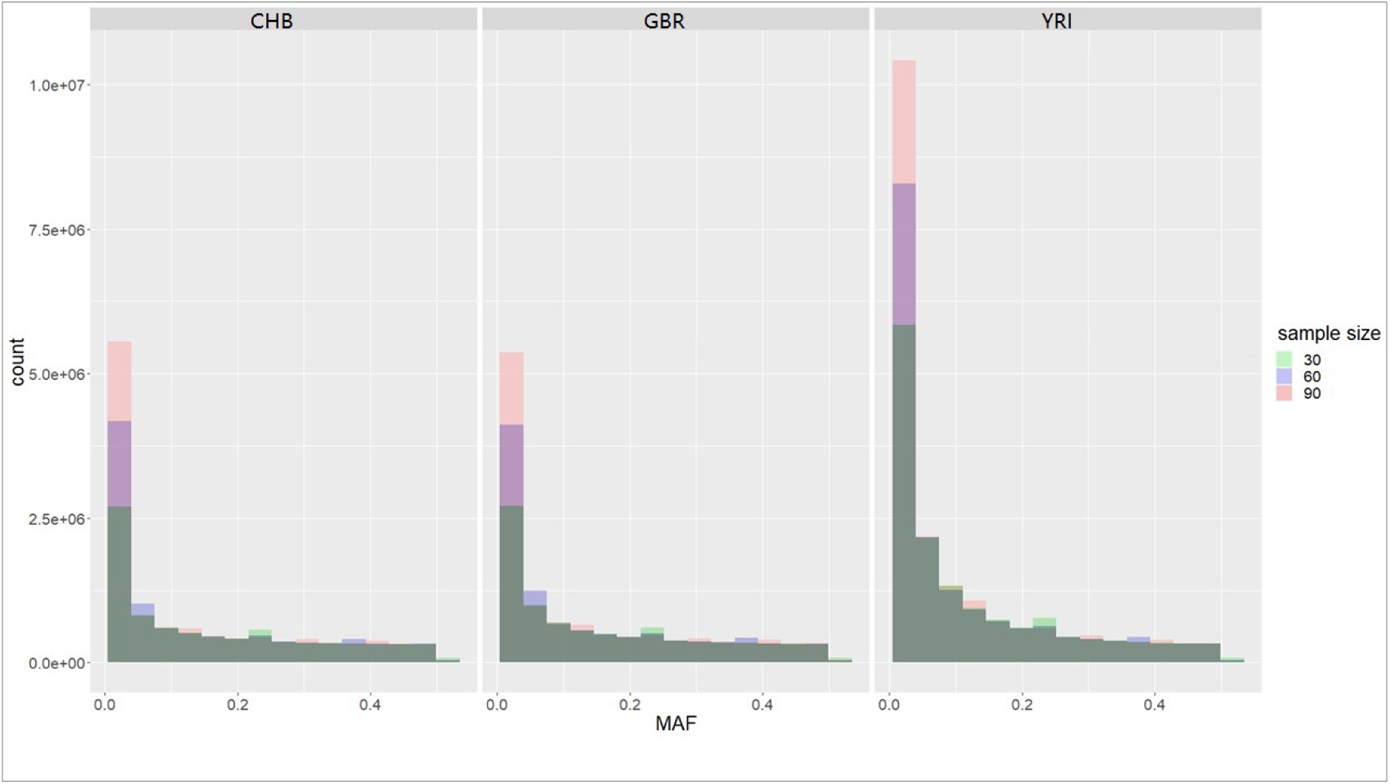
Distribution of MAF in reference data. Histograms present counts of MAF of subsets of referenced data. Light green shadow shows subsets of 30 referenced. Blue shows subsets of 60 referenced. Dusty rose shows subsets of 90 referenced. Dark green shows overlap of subsets of three sizes. Purple shows overlap between subsets of 30 and 60 referenced.

**Supplementary Figure 6:**
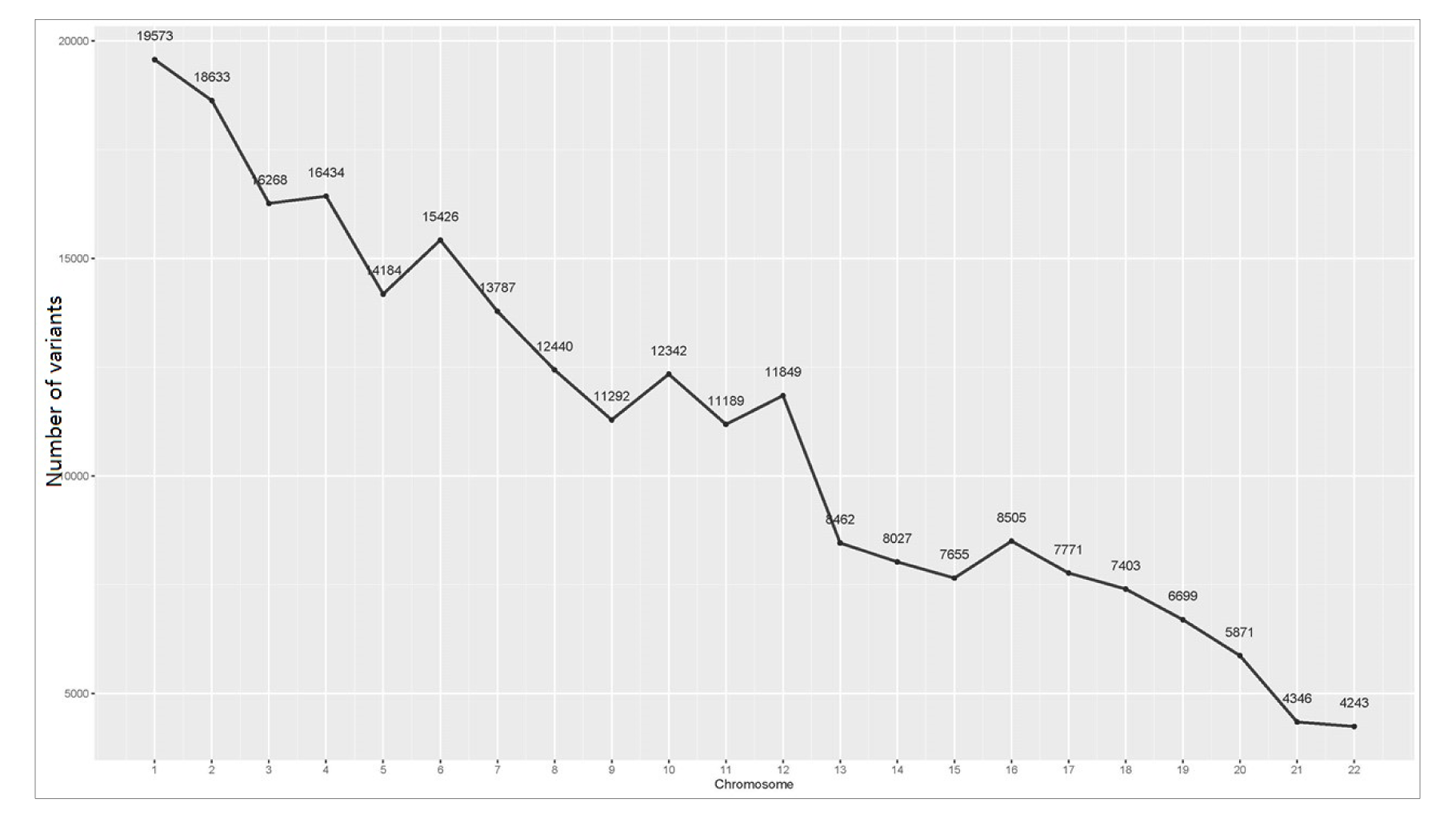
Number of variants pruned for PCA and kinship analysis, by chromosomes.

**Supplementary Figure 7:**
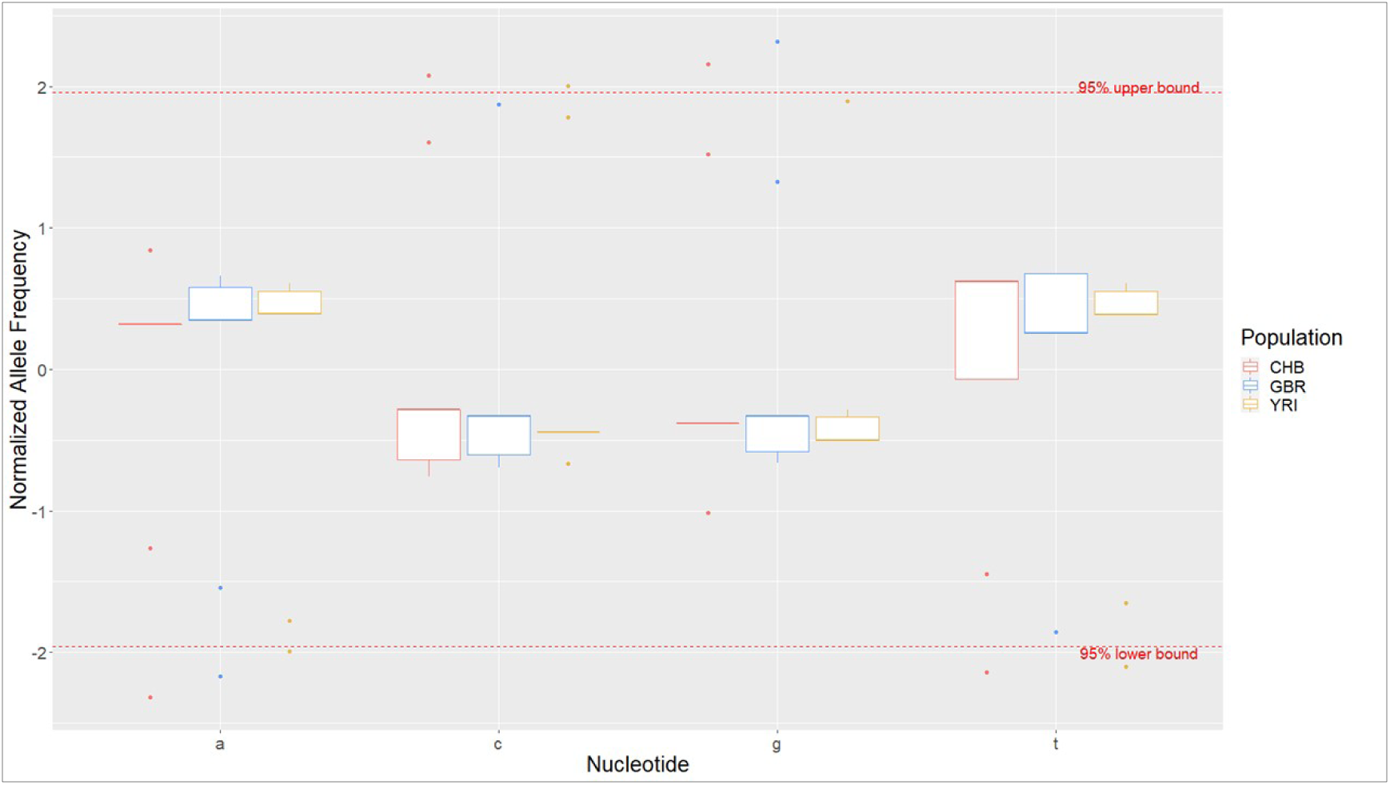
Boxplot of Post-simulation nucleotide frequency with largest potential of variance. CHB boxes and outliers are in red; GBR are in blue; YRI are yellow. Simulations considered largest potential of variance are using chromosome 22, an input sample size of 30 and 300 number of generation passes under a WF model. Dash lines are 95% confidence interval bounds.

**Supplementary Figure 8:**
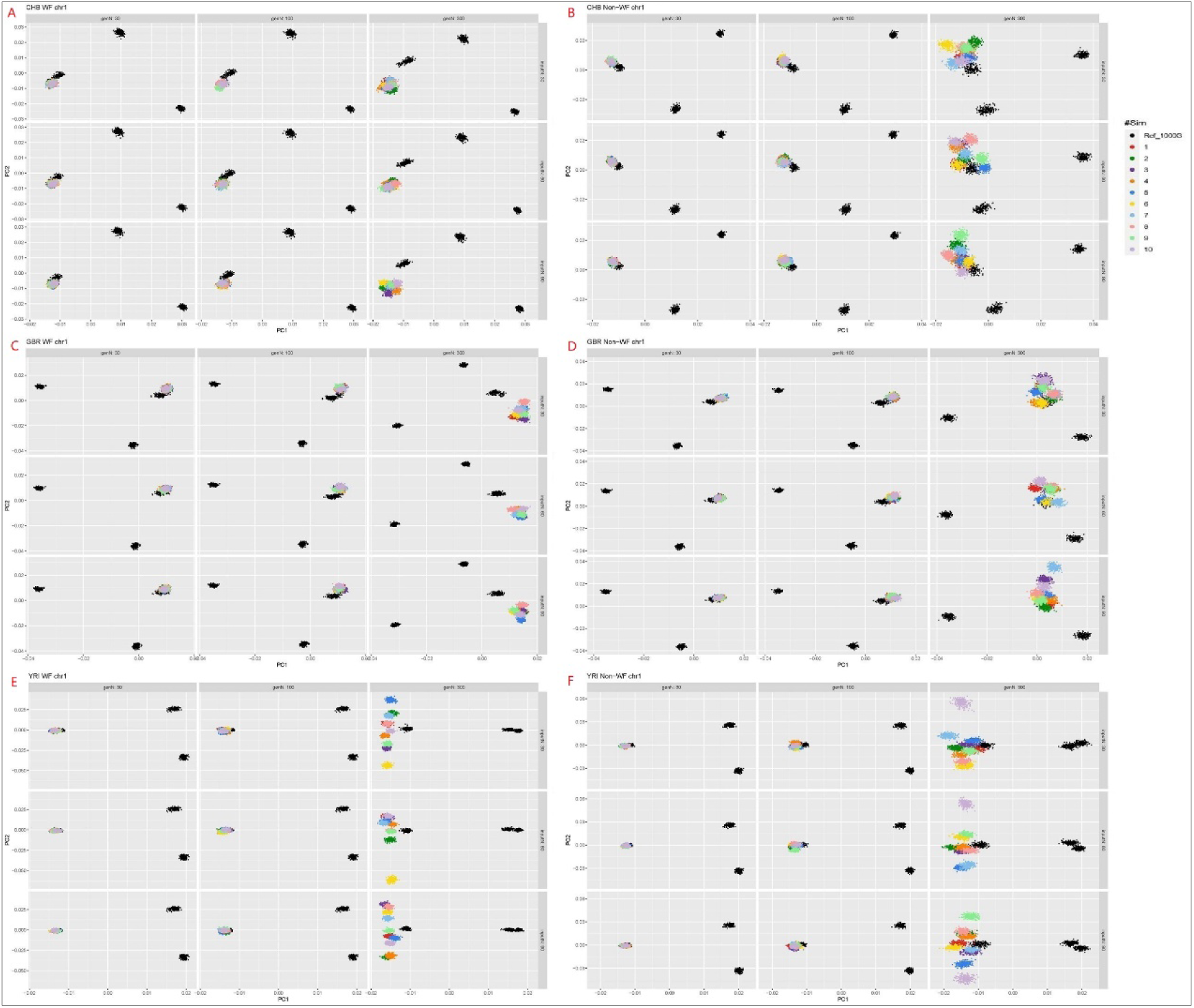
Scatter plots of PC1 vs PC2 of simulated data in chromosome one. Black dots are plots of references from 1KGP. Other colored dots are data simulated from ten parallel runs. **A**. CHB referenced and under WF model. **B**. CHB referenced and under Non-WF model. **C**. GBR referenced and under WF model. **D**. GBR referenced and under Non-WF model. **E**. YRI referenced and under WF model. **F**. YRI referenced and under Non-WF model.

**Supplementary Figure 9:**
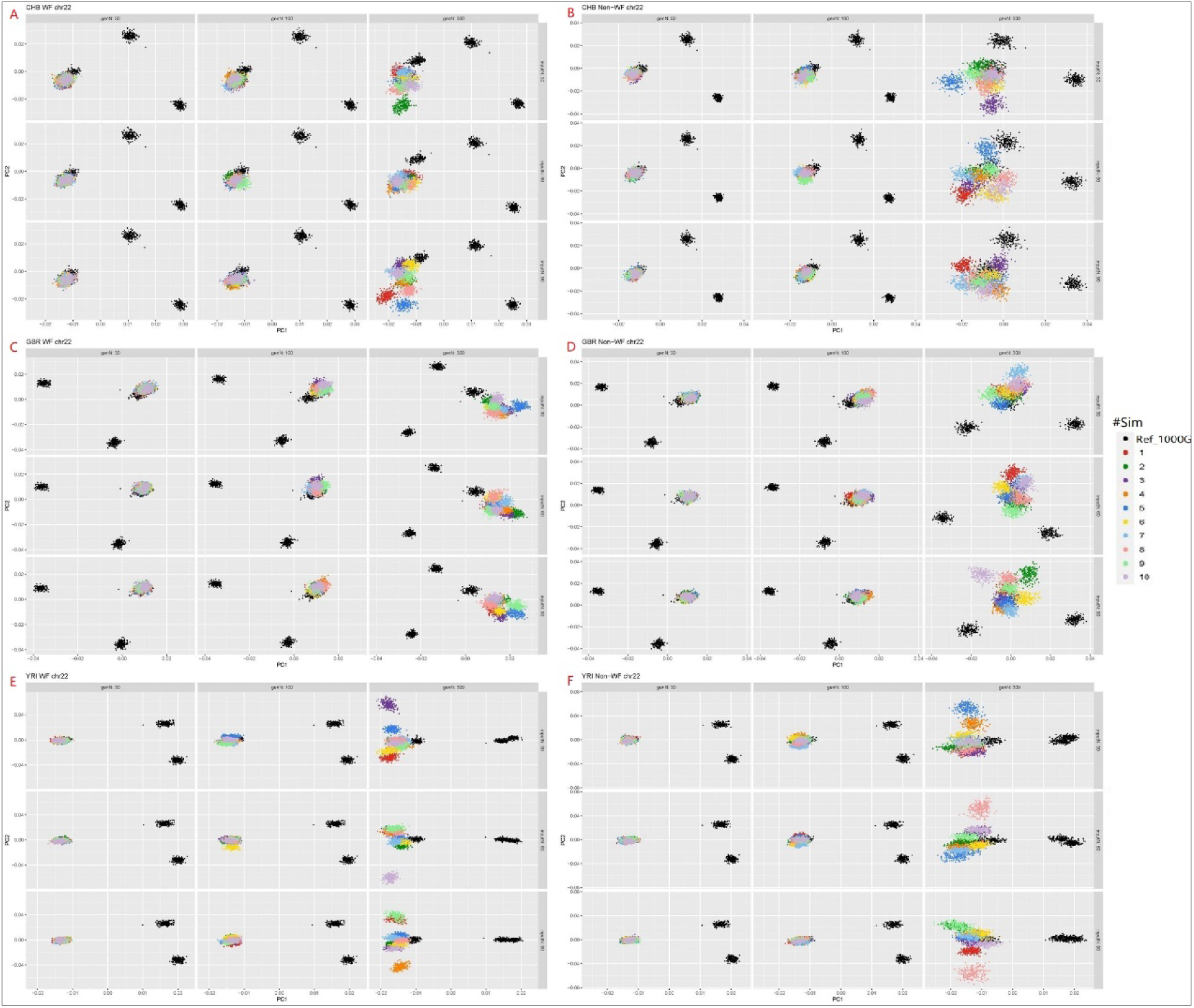
Scatter plots of PC1 vs PC2 of simulated data in chromosome 22. Black dots are plots of references from 1KGP. Other colored dots are data simulated from ten parallel runs. **A**. CHB referenced and under WF model. **B**. CHB referenced and under Non-WF model. **C**. GBR referenced and under WF model. **D**. GBR referenced and under Non-WF model. **E**. YRI referenced and under WF model. **F**. YRI referenced and under Non-WF model.

**Supplementary Figure 10:**
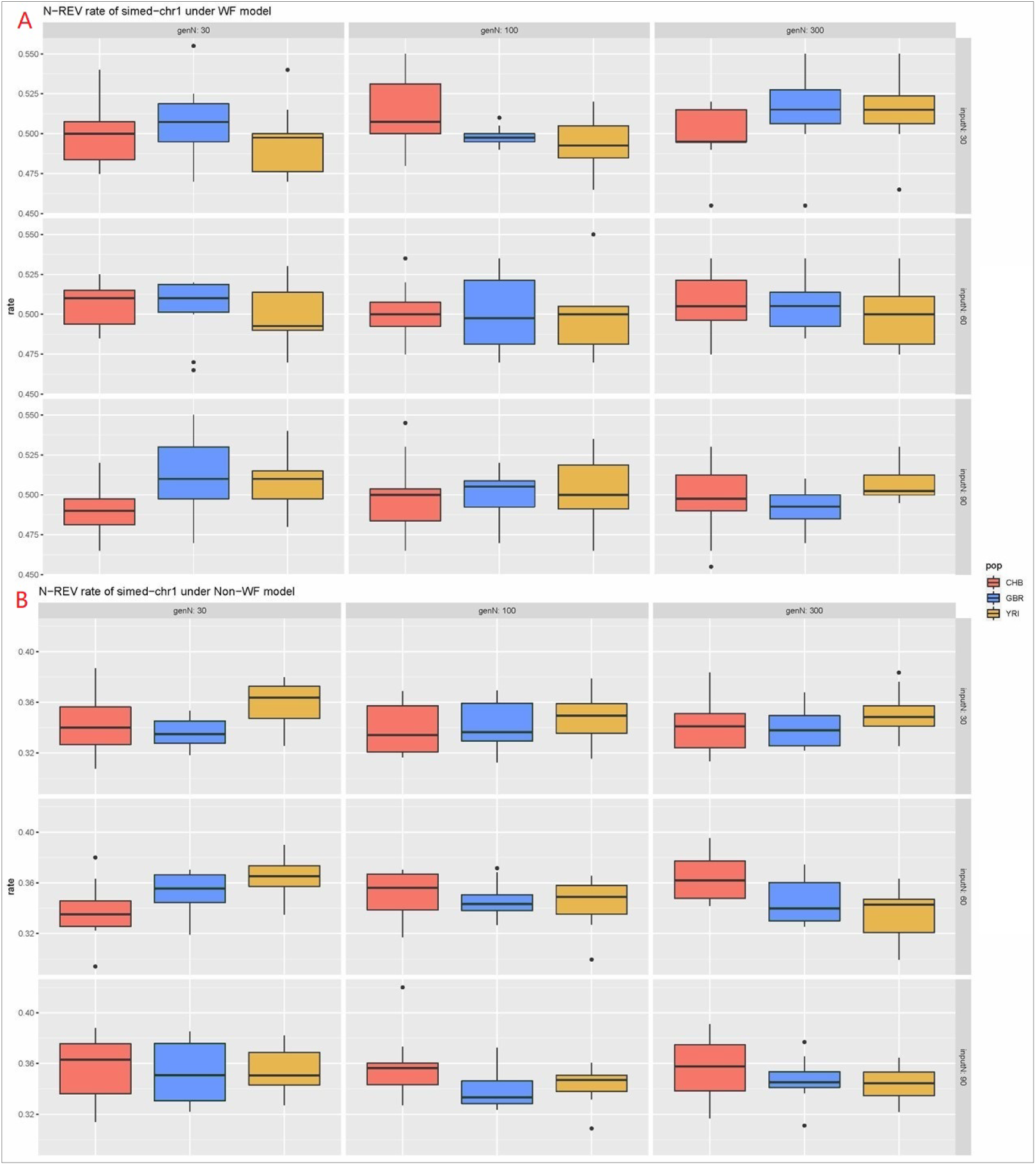
Box plots of N-REV rate estimates of simulated data in chromosome one. CHB in red; GBR in blue; YRI in yellow. Data are from simulations varying number of generation passes (30,100,300) by columns and input sample sizes (30,60,90) by rows. **A**. Under WF model. **B**. Under Non-WF model.

**Supplementary Figure 11:**
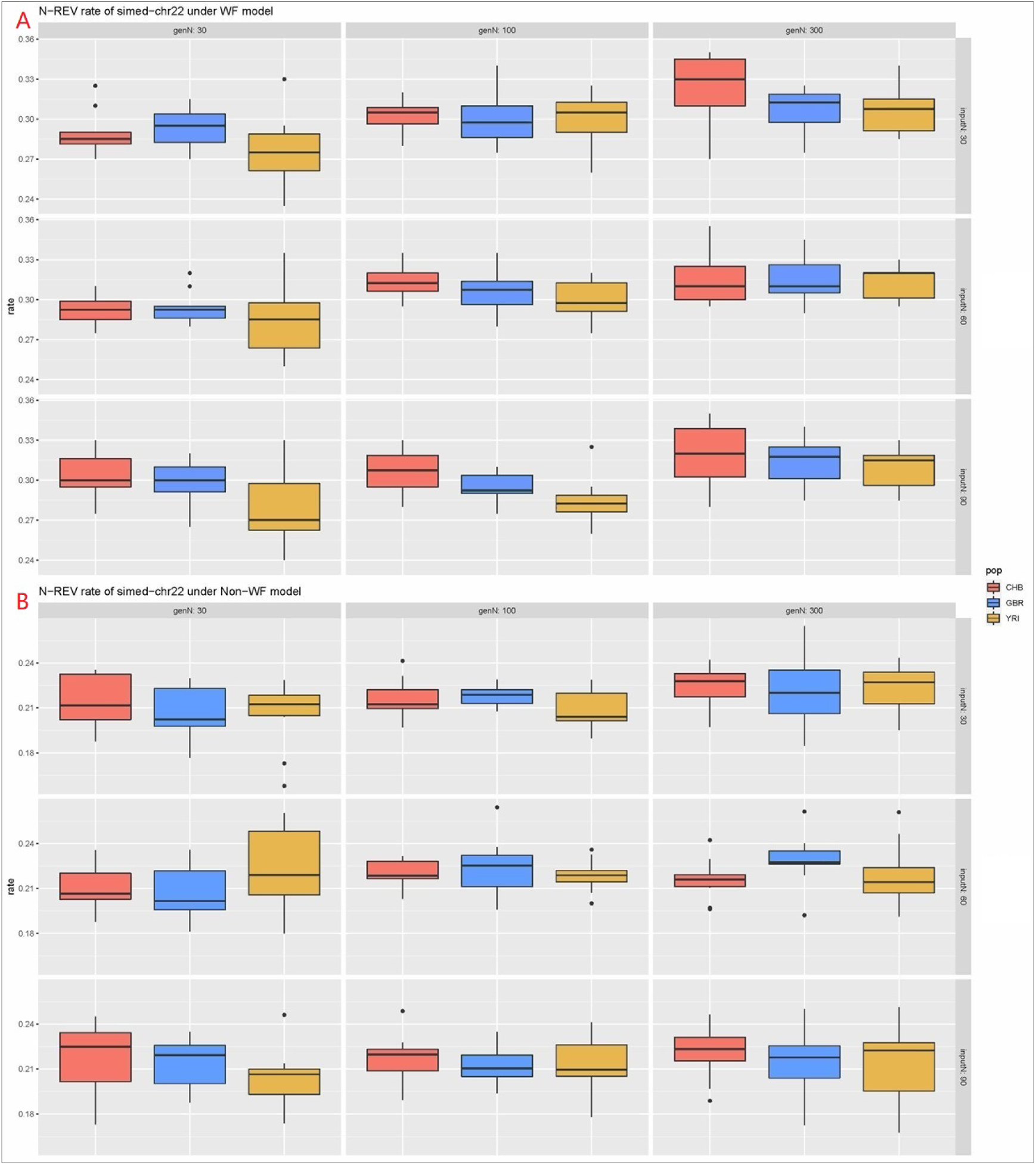
Box plots of N-REV rate estimates of simulated data in chromosome 22. CHB in red; GBR in blue; YRI in yellow. Data are from simulations varying number of generation passes (30,100,300) by columns and input sample sizes (30,60,90) by rows. **A**. Under WF model. **B**. Under Non-WF model.

**Supplementary Figure 12:**
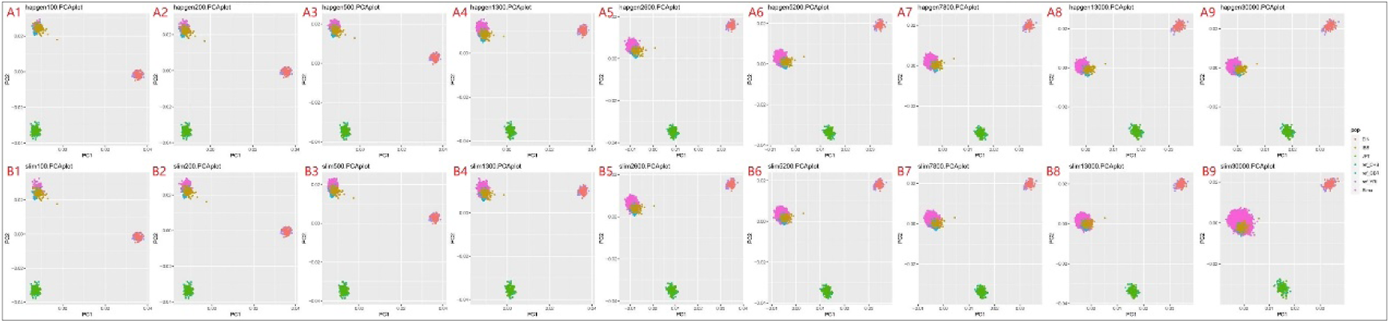
Comparison of PCA plots of data in chromosome 15 from SLiM and HapGen simulations. Plots of simulated data colored in pink. Input sample size = 90. Number of generation passes = 30. **A**. Use HapGen for simulation. **B**. Use SLiM for simulation. **1**. N (output sample size) = 100. **2**. N = 200. **3**. N = 500. **4**. N = 1300. **5**. N = 2600. **6**. N = 5200. **7**. N = 7800. **8**. N = 13000. **9**. N = 30000.

**Supplementary Figure 13:**
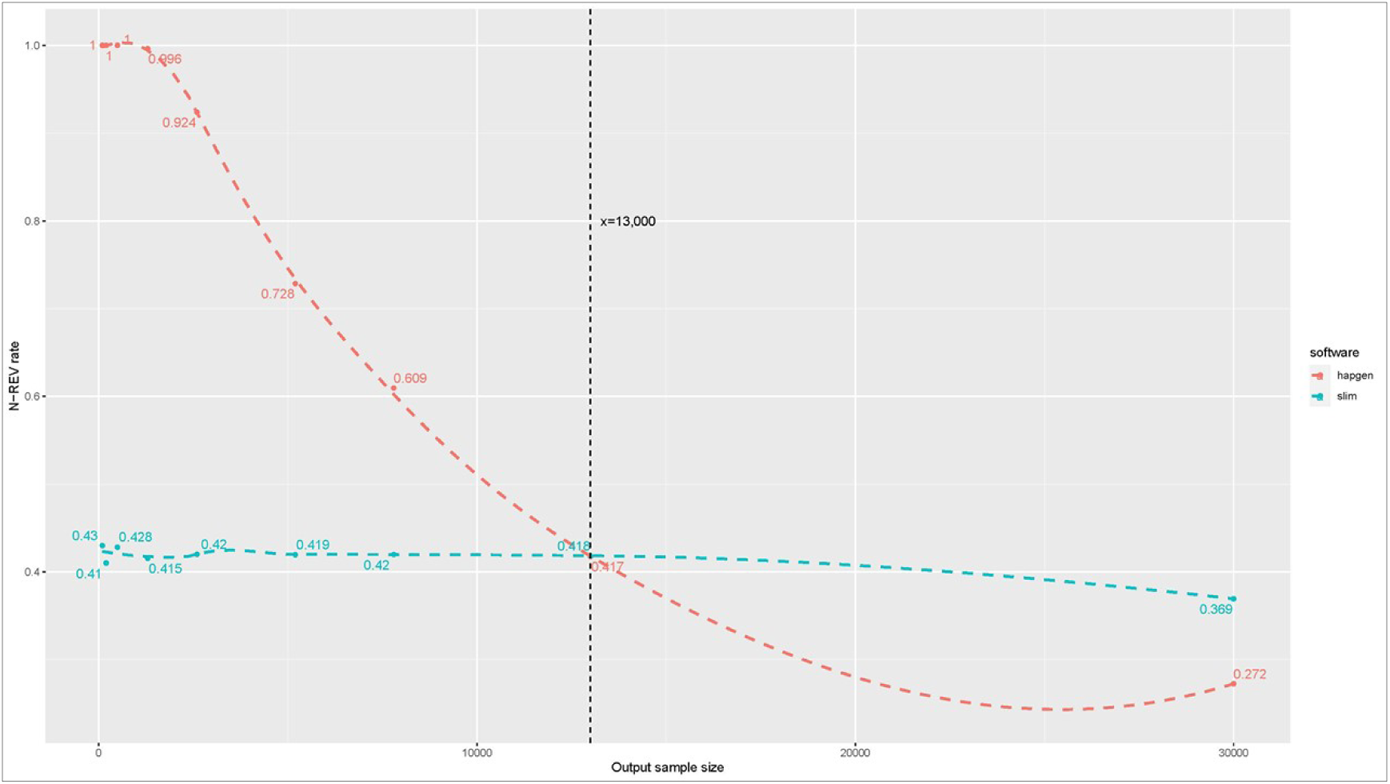
Scatter plots of N-REV rates vs output sample sizes comparing simulations using SLiM and HapGen. Plots of simulated data in chromosome 15. Input sample size = 90. Number of generation passes = 30. Y-axis of Colored dots are estimated N-REV rates and X-axis are sample sizes. Simulation using HapGen colored in red orange. Simulation using SLiM colored in Turkish green. Dash lines are smooth lines based on Locally Weighted Least Squares Regression (LOESS)

**Supplementary Figure 14:**
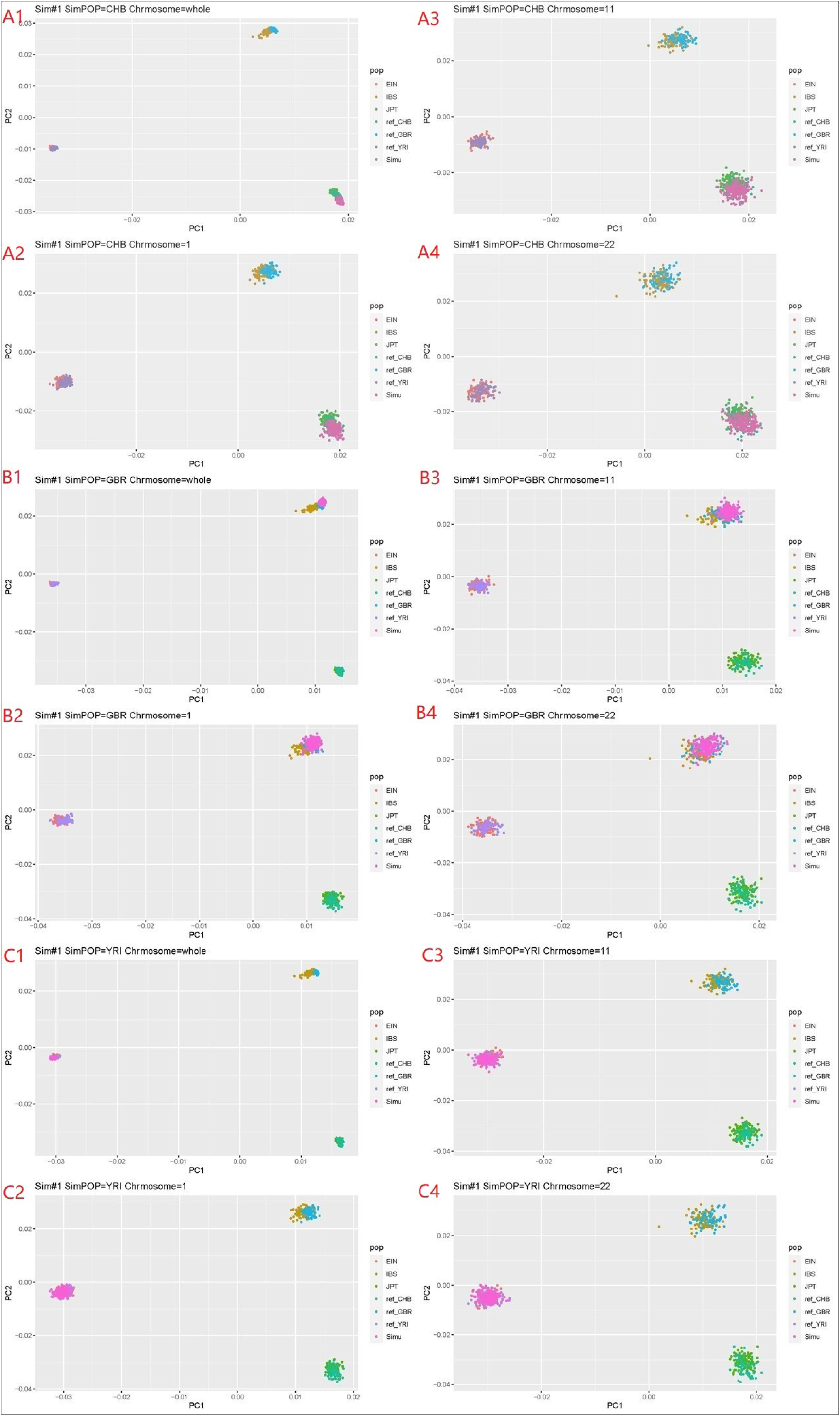
Plots of PC1 vs PC2 from whole-genome simulations, derived from the genome-wide and chromosome-level. Plots of simulated data colored in pink. Input sample size = 60. Output sample size = 200. Number of generation passes = 30. **A**. CHB referenced. **B**. GBR referenced. **C**. YRI referenced. **1**. PCA derived from whole-genome pruned data. **2**. PCA derived from pruned data in chromosome 1. **3**. PCA derived from pruned data in chromosome 11. **4**. PCA derived from pruned data in chromosome 22.

## Notes

### Competing Interest Statement

The authors have declared no competing interest.

